# The chromosomal distribution of sex-biased microRNAs in *Drosophila* is non-adaptive

**DOI:** 10.1101/2021.09.25.461796

**Authors:** Antonio Marco

## Abstract

Genes are often differentially expressed between males and females. In *Drosophila melanogaster*, the analysis of sex-biased microRNAs (short non-coding regulatory molecules) has revealed striking differences with protein-coding genes. Mainly, the X chromosome is enriched in male-biased microRNA genes, although it is depleted of male-biased protein-coding genes. The paucity of male-biased genes in the X chromosome is generally explained by an evolutionary process called demasculinization. I suggest that the excess of male-biased microRNAs in the X chromosome is due to high-rates of de novo emergence of microRNAs (mostly in other neighboring microRNAs), a tendency of novel microRNAs in the X chromosome to be expressed in testis, and to a lack of a demasculinization process. To test this hypothesis I analysed the expression profile of microRNAs in males, females and gonads in *D. pseudoobscura*, in which an autosome translocated into the X chromosome effectively becoming part of a sex chromosome (neo-X). I found that the pattern of sex-biased expression is generally conserved between *D. melanogaster* and *D. pseudoobscura*. Also, orthologous microRNAs in both species conserve their chromosomal location, indicating that there is no evidence of demasculinization or other inter-chromosomal movement of microRNAs. *D. pseudoobscura*specific microRNAs in the neo-X chromosome tend to be male-biased and particularly expressed in testis. In summary, the apparent paradox resulting from male-biased protein-coding genes depleted in the X chromosome and an enrichment in male-biased microRNAs is consistent with different evolutionary dynamics between coding genes and short RNAs.

## INTRODUCTION

Gene expression is tightly regulated by mechanisms that make gene products to be expressed in specific organs at specific times. This spatiotemporal control of gene expression determines the development of fertilized eggs into adult organisms (Davidson 2006; Wolpert et al. 2006). In organisms with two sexes a large fraction of genes are also differentially expressed between males and females (Parisi et al. 2003; Ranz et al. 2003). Also, most male or female specific expression occurs in the gonads, alhough other tissues also show sex-expression bias (Parisi et al. 2004; Chang et al. 2011). Female biased genes tend to be located in the X chromosome (in XY systems) and male-biased genes tend to be depleted in the X chromosome (Parisi et al. 2003; Ranz et al. 2003; Khil et al. 2004).

In the model species *Drosophila melanogaster* the study of sex-biased expression indicates that male-biased genes often appear in the X chromosome (Arguello et al. 2006; Levine et al. 2006; Begun et al. 2007; Chen et al. 2007), but they are later retroposed (copied) to an autosome and the original copy is eventually lost (Betran et al. 2002; Zhang, Vibranovski, Krinsky, et al. 2010). This process, called ‘demasculinization’, was predicted already by early theoretical models (Rice 1984), and recent works seems to support it (Jiang and Machado 2009; Bachtrog et al. 2010; Zhang, Vibranovski, Landback, et al. 2010; Zhang, Vibranovski, Krinsky, et al. 2010), although there is still some controversy around it, and alternative models should be taken into consideration (Meiklejohn and Presgraves 2012). However, these conclusions cannot be extrapolated to all gene products, as most work on sex-biased expression has been focused on a specific type of gene: protein-coding genes. Characterising the distinct evolutionary dynamics of different types of genes is paramount in evolutionary biology, as it can reveal how genetic and genomic features influence, and ultimately determine, the fate of genes (Lynch 2007).

MicroRNAs, short RNA post-transcriptional regulators [reviewed in (Bartel 2004; Axtell et al. 2011; Marco, Ninova, and Griffiths-Jones 2013)], are also expressed differently in males and in females (Marco, Kozomara, et al. 2013; Marco 2014; Marco 2015; Warnefors et al. 2017; Fowler et al. 2019). Although an early investigation suggested that they were also subject to demasculinization (Zhang, Vibranovski, Krinsky, et al. 2010), a more recent work indicates that this is probably not the case (Marco 2014). Indeed, male microRNAs tend to be enriched in the X chromosome (Mishima et al. 2008; Li et al. 2010), contrary to what is observed in proteincoding genes. In *D. melanogaster* novel microRNAs in the X chromosome are often expressed in testis and most are evolutionarily young (Marco 2014). To investigate the evolutionary dynamics of microRNAs in the X chromosome I characterised the sex-biased microRNA complement of the species *Drosophila pseudoobcura*, which diverged from *D. melanogaster* ~37 Myr ago (Hedges et al. 2015), and had an autosome translocated into an X-chromosome, becoming a newly evolved X-chromosome (or neo-X). This neo-X emerged about 13 Mys ago (Kaiser and Bachtrog 2010) which, despite being a short evolutionary period, has been enough to identify demasculinization affecting protein-coding genes in several studies (Zhang, Vibranovski, Krinsky, et al. 2010; Nozawa et al. 2014). Indeed, it has been estimated that, in *Drosophila*, one retrogene emerges every 2 Myr years at an approximate constant rate (Bai et al. 2007), and we would expect (at least in theory) a similar rate for microRNA genes. Hence, this system will shed light on the evolution of chromosomal composition of sex-biased microRNAs.

## RESULTS

In order to characterize differentially expressed microRNAs between males and females in *D. pseudoobscura* I analyzed three available small RNA datasets for *D. pseudoobscura*, two male samples and one female sample (Mohammed et al. 2018). However, these datasets have either no biological replicates, or they have been sequencing using different library construction methods. Hence, I also sequenced two male and two unfertilised female samples in two paired sequencing reactions (one male sample and one female sample in each sequencing array) using the same library preparation protocols. The differential gene expression analysis for this set was performed taking into account batch (paired) effects. For simplicity, the first set is called in this paper the *unpaired* dataset, and the newly reported here the *paired* dataset.

The differential expression analysis of the *unpaired* dataset, for a False Discovery Rate (FDR) or 10% and a expression difference of 25% or over, identified 38 microRNAs with sex-biased expression: 19 overexpressed in females and 19 overexpressed in males (Figure 1A, Supplementary Table 1). As expected (see Discussion) the *paired* experiments identified less biased microRNAs: 12 overexpressed in females and 10 in males (Figure 1B, Supplementary Table 1). For a more stringent FDR threshold of 5%, the unpaired and paired experiments identified 25 and 15 differentially expressed microRNAs respectively. The code available in GitHub allows the user to modify the different selection criteria. For the dataset with replicates in both conditions (paired dataset), an alternative method based on read count transformation and empirical Bayes yielded comparable results: 19 overexpressed in females and 10 overexpressed in males (Supplementary Figure 1). Although there are some differences in the significance values associated with individual microRNAs, the fold change estimation (which is used in the subsequent analyses) for both methods was consistent (Supplementary Figure 1).

**Figure 1.**
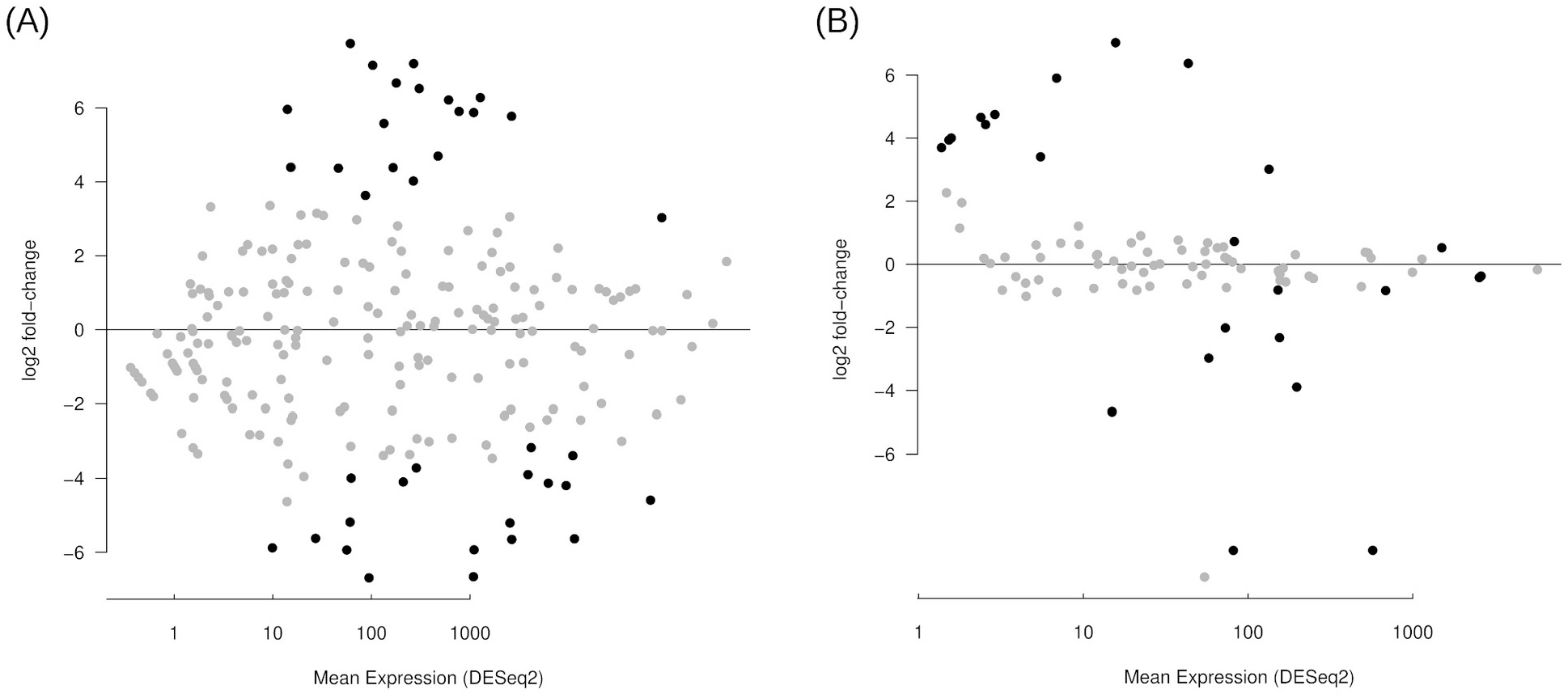
Sex-biased expression of microRNAs in *Drosophila pseudoobscura*. Smear plots of expressed microRNAs. Black dots are microRNAs with a significant sex-biased expression (see Methods). Positive values indicate male-biased microRNA expression, and negative values female-biased microRNA expression. (A) Expression profile for the unpaired dataset. (B) Expression profile for the paired dataset.

To investigate whether the chromosomal location between *D. melanogaster* and *D. pseudoobscura* orthologs is conserved, the one-to-one orthologous microRNAs as annotated in (Mohammed et al. 2018) were first considered. There were 128 annotated orthologs, of which 122 were successfully mapped to the latest genome versions. The chromosome in which each microRNA is located is conserved for all studied sequences, although some evidence of translocations and inversion are observed within chromosomes (Figure 2). Importantly, among this set of orthologs, there is not a single case of a microRNA in one of the *Drosophila* species that moved to a different chromosome in the other species. In other words, this specific analysis did not identify a microRNA that moved out of a sex chromosome to an autosome (or vice versa), a phenomena that is well described for protein-coding genes, and it is generally due to retroposition ((Sturgill et al. 2007); see Discussion).

**Figure 2.**
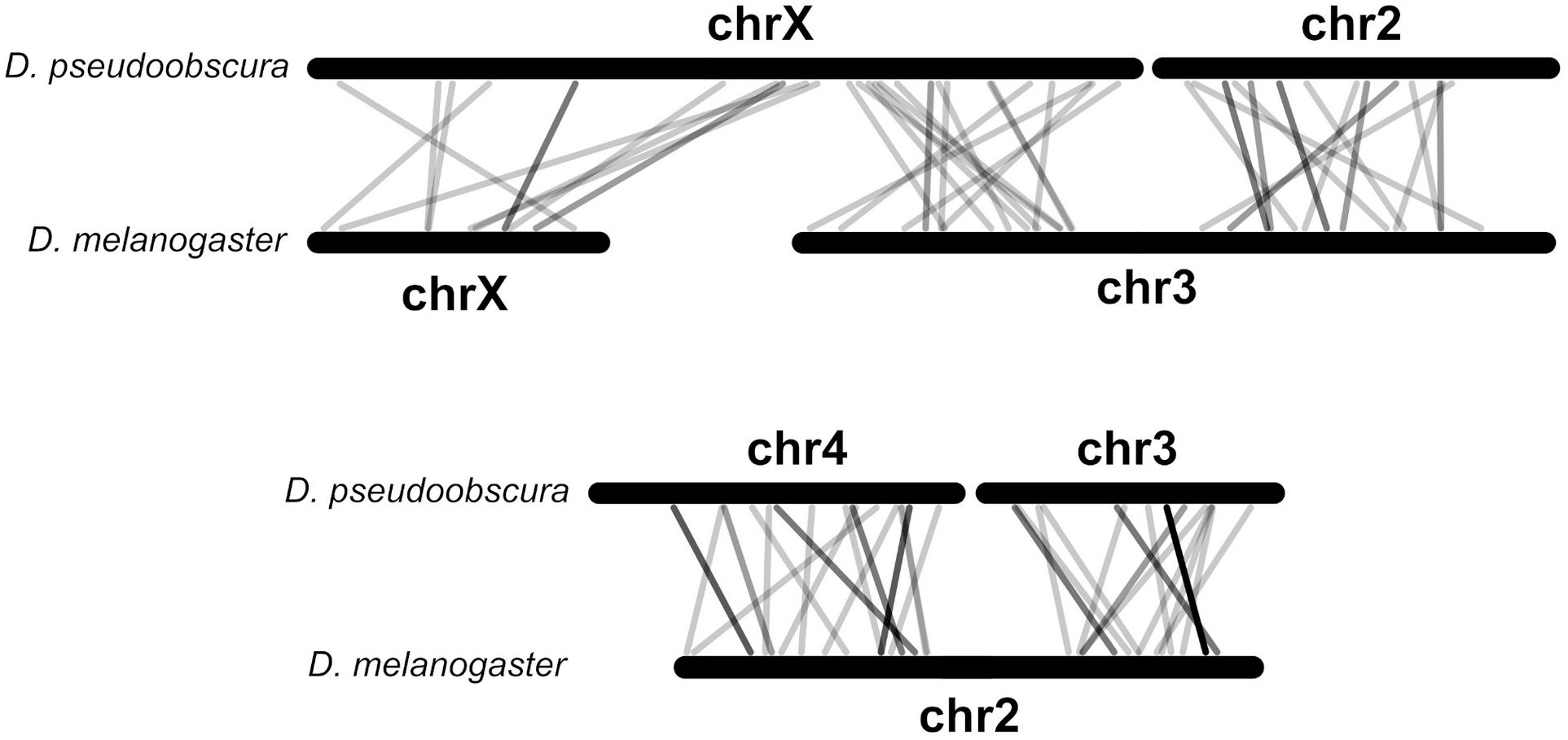
Comparison of the chromosomal location of microRNAs between *D. melanogaster* and *D. pseudoobscura*. Homologous chromosomes of both species are paired, and grey lines connect the location of orthologous microRNAs. Dark grey lines indicate overlap of two or more lines, the darker the more microRNAs: this is due to clustered microRNAs.

A potential bias in the chromosomal location analysis is that orthology may have been defined, partly, by using chromosomal context. To control for that I run reciprocal BLAST among all *D. melanogaster* and *D. pseudoobscura* annotated microRNAs (see Methods). Clusters of similar sequences were aligned and similarity trees were built (Supplementary File 1). Only one microRNA in *D. pseudoobscura* remained unpaired to an ortholog: dps-mir-92c. Mir-92 is one of the most conserved microRNA families in animals, and it has two copies in *D. melanogaster* (mir-92a and mir-92b) and three (mir-92a, mir-92b and mir-92c) in *D. pseudoobscura*. Recent work has shown that mir-92c is deeply conserved in insects and it is not associated with a recent duplication of any of the other mir-92 in the pseudoobscura lineage (Ninova et al. 2014), so it seems that it has been lost in *D. melanogaster*, although its homology relationships with other microRNA family members is not fully understood. As a matter of fact, Mohammed et al. (2018) suggest that *D. melanogaster* mir-311 is mir-92c’s ortholog, a relationship maintained in the analyses for consistency (see below; Figure 3), but that did not alter the results.

**Figure 3.**
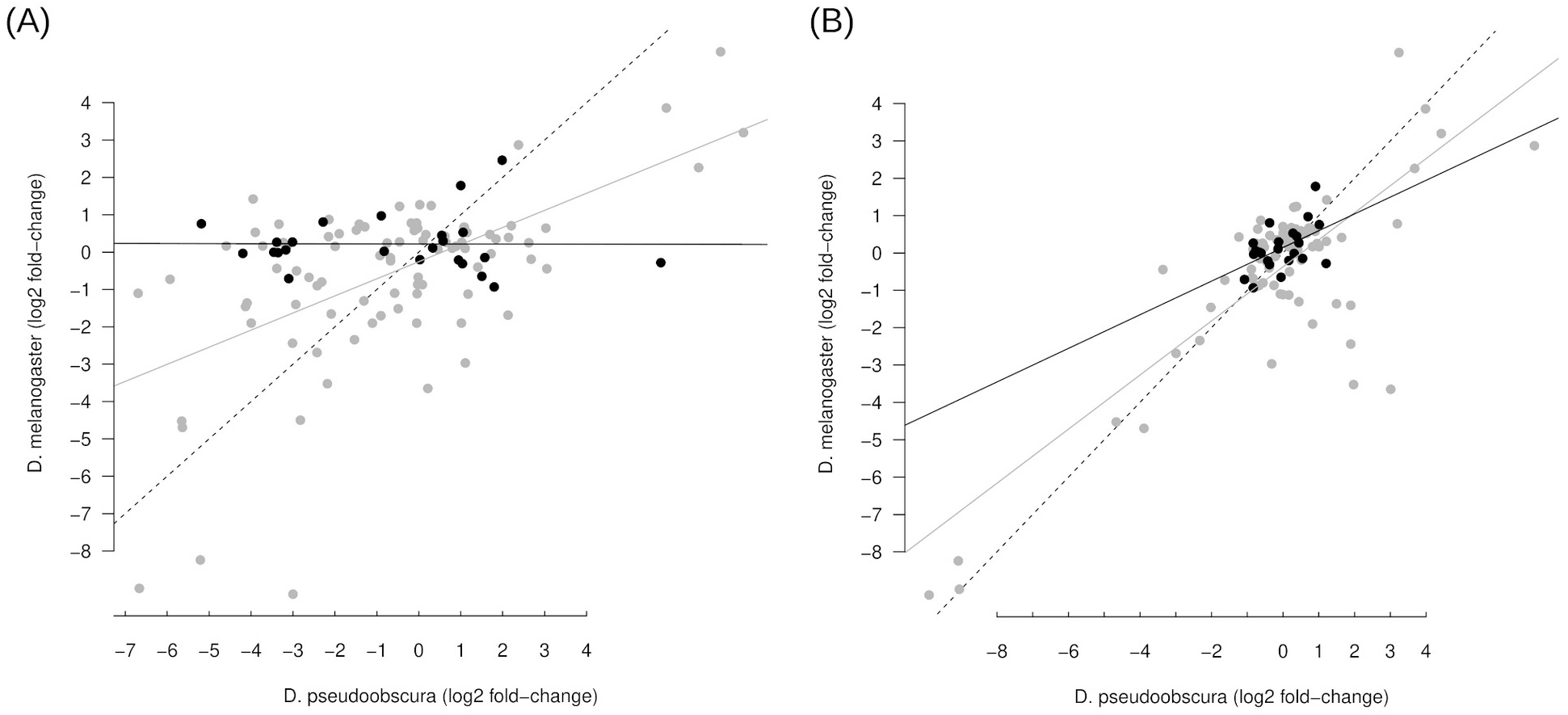
Conservation of sex-biased microRNA expression. Scatter-plot of log2 fold-change expression values of *D. melanogaster* and *D. pseudoobscura*. Black dots represent microRNAs located in the neo-X chromosome in *D. pseudoobscura*, and the other microRNAs are in grey, and black and grey straight lines represent the linear model fitted to both groups respectively. Dashed line is the 1:1 ratio. (A) Comparison between *D. melanogaster* and the unpaired dataset. (B) Comparison between *D. melanogaster* and the paired dataset.

In addition, the *D. pseudoobscura* genome was scanned for potential homologous microRNAs annotated so far only in *D. melanogaster* (see Methods), to identify copies that may have not been detected in earlier works (either because of limited genome sequence information or by the use of strict criteria to filter putative microRNAs). The BLAST searches predicted 26 putative one-to-one previously not identified microRNA orthologs between both species (Table 1). Only one newly identified microRNAs was in different chromosomes between the two species: dme-mir-2281. However, independently of whether this is a bona fide microRNA (it is lowly expressed and poorly conserved), it would be an autosome to autosome translocation. There were two microRNAs in *D. melanogaster* that have two copies in *D. pseudoobscura* with at least one of those not previously annotated (Table 2). In all these cases, all copies were in the analogous chromosomes in both species.

**Table 1.**
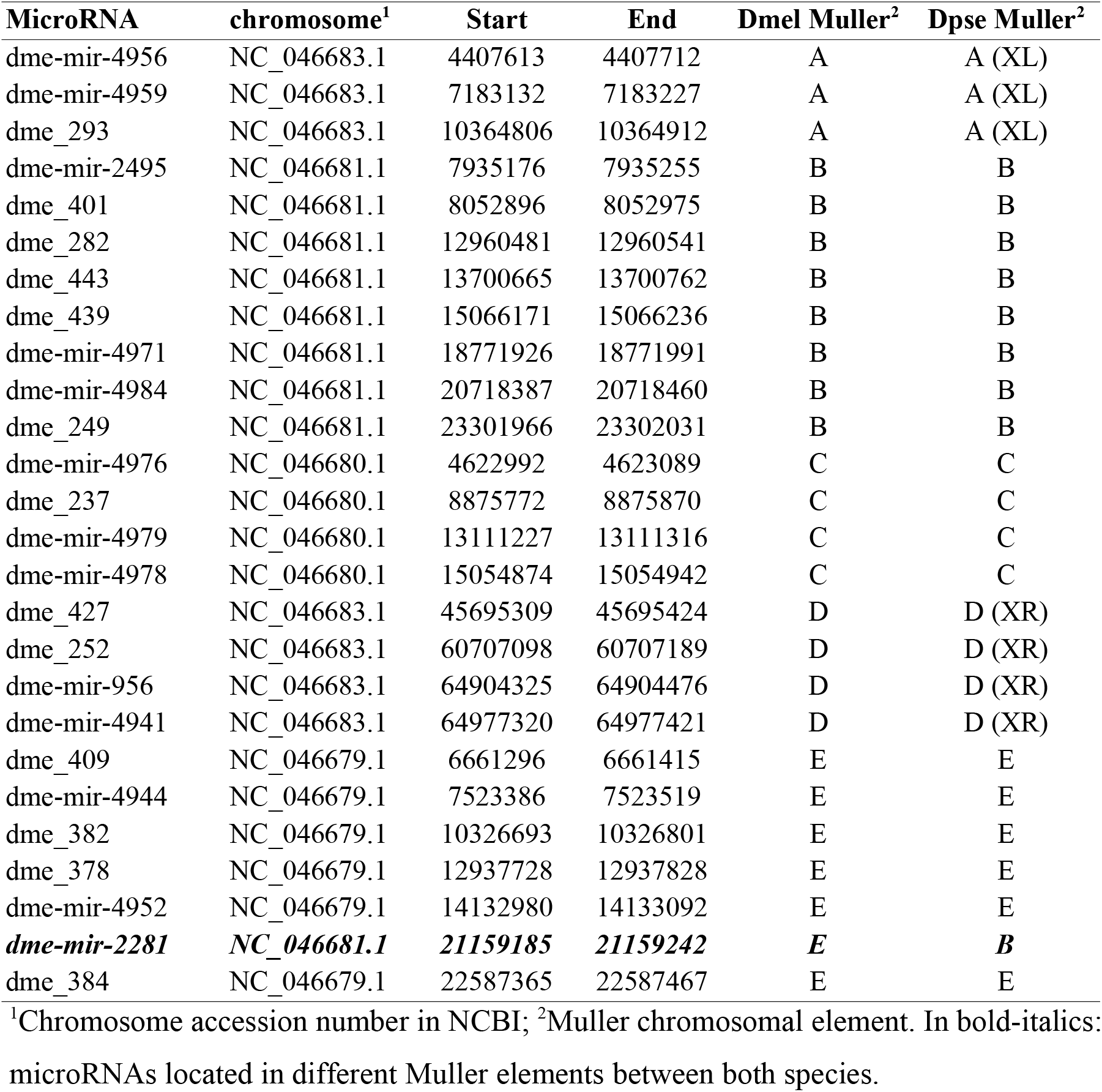
Predicted newly discovered *D. pseudoobscura* microRNAs from similarity analysis.

| MicroRNA | chromosome<sup>1</sup> | Start | End | Dmel Muller<sup>2</sup> | Dpse Muller<sup>2</sup> |
| --- | --- | --- | --- | --- | --- |
| dme-mir-4956 | NC_046683.1 | 4407613 | 4407712 | A | A (XL) |
| dme-mir-4959 | NC_046683.1 | 7183132 | 7183227 | A | A (XL) |
| dme_293 | NC_046683.1 | 10364806 | 10364912 | A | A (XL) |
| dme-mir-2495 | NC_046681.1 | 7935176 | 7935255 | B | B |
| dme_401 | NC_046681.1 | 8052896 | 8052975 | B | B |
| dme_282 | NC_046681.1 | 12960481 | 12960541 | B | B |
| dme_443 | NC_046681.1 | 13700665 | 13700762 | B | B |
| dme_439 | NC_046681.1 | 15066171 | 15066236 | B | B |
| dme-mir-4971 | NC_046681.1 | 18771926 | 18771991 | B | B |
| dme-mir-4984 | NC_046681.1 | 20718387 | 20718460 | B | B |
| dme_249 | NC_046681.1 | 23301966 | 23302031 | B | B |
| dme-mir-4976 | NC_046680.1 | 4622992 | 4623089 | C | C |
| dme_237 | NC_046680.1 | 8875772 | 8875870 | C | C |
| dme-mir-4979 | NC_046680.1 | 13111227 | 13111316 | C | C |
| dme-mir-4978 | NC_046680.1 | 15054874 | 15054942 | C | C |
| dme_427 | NC_046683.1 | 45695309 | 45695424 | D | D (XR) |
| dme_252 | NC_046683.1 | 60707098 | 60707189 | D | D (XR) |
| dme-mir-956 | NC_046683.1 | 64904325 | 64904476 | D | D (XR) |
| dme-mir-4941 | NC_046683.1 | 64977320 | 64977421 | D | D (XR) |
| dme_409 | NC_046679.1 | 6661296 | 6661415 | E | E |
| dme-mir-4944 | NC_046679.1 | 7523386 | 7523519 | E | E |
| dme_382 | NC_046679.1 | 10326693 | 10326801 | E | E |
| dme_378 | NC_046679.1 | 12937728 | 12937828 | E | E |
| dme-mir-4952 | NC_046679.1 | 14132980 | 14133092 | E | E |
| ***dme-mir-2281*** | ***NC_046681.1*** | ***21159185*** | ***21159242*** | ***E*** | ***B*** |
| dme_384 | NC_046679.1 | 22587365 | 22587467 | E | E |

<sup>1</sup>Chromosome accession number in NCBI; <sup>2</sup>Muller chromosomal element. In bold-italics: microRNAs located in different Muller elements between both species.

**Table 2.** Predicted newly discovered *D. pseudoobscura* with two copies.

| **MicroRNA** | **chromosome<sup>1</sup>** | **Start** | **End** | **Dmel Muller<sup>2</sup>** | **Dpse Muller<sup>2</sup>** |
| --- | --- | --- | --- | --- | --- |
| dme_393 | NC_046679.1 | 21527631 | 21527726 | E | E |
| dme_393 | NC_046679.1 | 21530842 | 21530915 | E | E |
| dme-mir-4948 | NC_046679.1 | 1526563 | 1526662 | E | E |
| dme-mir-4948 | NC_046679.1 | 2671493 | 2671578 | E | E |

<sup>1</sup>Chromosome accession number in NCBI; <sup>2</sup>Muller chromosomal element. In bold-italics: microRNAs located in different Muller elements between both species.

Finally, I considered those *D. melanogaster* microRNAs with more than two copies in the *D. pseudoobscura* genome. There were 6 microRNAs in this category (dme_208, dme_417, dme_455, dme_5, dme_7 and dme-mir-10404), all hitting 55 positions in the genome. These sequences are actually derived from rRNAs, and similar sequences are found associated to rRNA genes in other insects (Chak et al. 2015). In summary, these comparative genomics analyses did not reveal any microRNA that may have moved out, or copied from, the X to any other autosomal chromosome.

To investigate whether the expression profile of microRNAs change in a different chromosomal context, I compared the sex-bias in expression of orthologogous microRNAs between both *Drosophila* species. A scatter plot of the log2 fold-change values for pairs of autosomal orthologs reveals that the expression bias is conserved between both species in the unpaired dataset, although microRNA expression biases in the *D. pseudoobscura* neo-X chromosome does not show any association with their ortholog expression biases (Figure 3A, Supplementary Table 1). In the paired dataset, where two biological replicates for both sexes exist and batch effects are controlled, the female/male expression bias is also roughly conserved in microRNAs that are autosomal in *D. melanogaster* but that are in the new-X chromosome in *D. pseudoobscura* (Figure 3B; fitted linear model p = 0.06 R^2^=0.2). There are, however, two notable exceptions of microRNAs that are strongly male biased in *D. pseudoobscura* but female biased in *D. melanogaster* (Figure 1B): mir-311 and mir-92c (see Discussion).

To characterize the expression of novel and conserved microRNAs depending on the chromosomal context, I compared the expression bias (log2 fold-change) of novel and conserved microRNAs in both autosomes and the X chromosome in *D. pseudoobscura*. By building a linear model on ranked log2 fold-changes considering chromosomal context and evolutionary age (Scheirer-Ray-Hare test), there was strong statistical support for novel microRNAs to be overexpressed in males, in both the unpaired (p=0.0045; Figure 4A) and the paired (p=0.0187; Figure 4B) datasets. In the controlled paired dataset, within the novel microRNAs, those in the X chromosome tend to have a higher expression bias towards males with respect to those in the autosomes (Figure 4B). More specifically, when I compare the testis to ovary expression ratio for microRNAs expressed in the gonads, a clear pattern of novel microRNAs in the X tending to be overexpressed in testis becomes evident (Figure 4C; conservation p<0.0001, chromosome p=0.0115).

**Figure 4.**
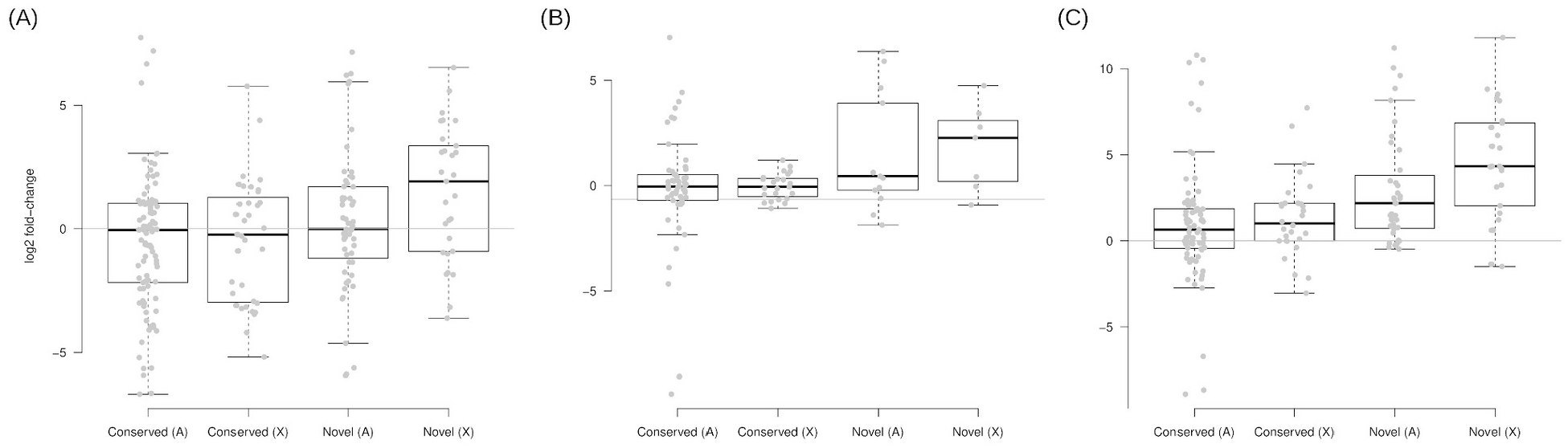
Expression of novel and conserved microRNAs. Log2 fold-change of male versus female expression for conserved or *D. pseudoobscura* specific (novel) microRNAs, either autosomal (A) or in the X chromosome (X) for the unpaired (A) and the paired (B) dataset. (C) Log2 fold-change of testis versus ovary expression.

A caveat in this analysis is the non-independence of expression profiles among clustered microRNAs, as they tend to be produced from the same transcript. To evaluate the impact of microRNA clusters, only the highest expressed microRNA in each cluster (microRNAs within 10Kb to each other) was considered. Overall the log2 fold change in expression is similar in both species (Figure 5A) with the main exception of the microRNA cluster already mentioned above. For novel X-linked microRNA clusters, there were four loci, all highly expressed in testis (Figure 5B). This also suggests that novel X-linked microRNAs tend to emerge within other microRNA clusters/loci (see Discussion).

**Figure 5.**
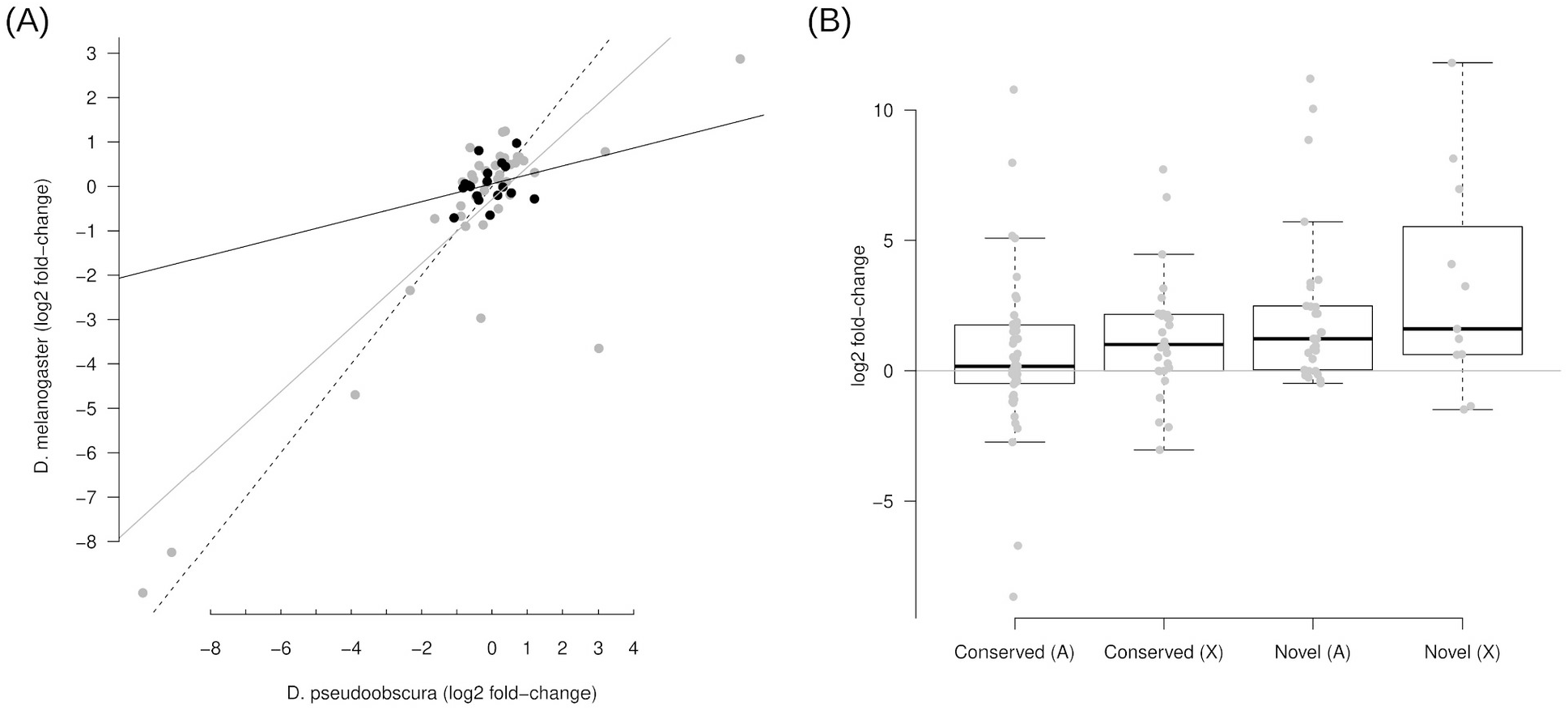
Sex-biased expression of microRNA clusters in *Drosophila pseudoobscura*. (A) Scatter-plot of log2 fold-change expression values of *D. melanogaster* and *D. pseudoobscura* clusters for the paired dataset. Black dots represent microRNA clusters located in the neo-X chromosome in *D. pseudoobscura*, and the other microRNAs are in grey, and black and grey straight lines represent the linear model fitted to both groups respectively. Dashed line is the 1:1 ratio. (B) Log2 fold-change of testis versus ovary expression for conserved or *D. pseudoobscura* specific (novel) microRNA clusters, either autosomal (A) or in the X chromosome (X).

## DISCUSSION

In this work I characterised the sex expression pattern of microRNAs in *Drosophila pseudoobscura* to investigate the evolutionary dynamics of male expressed microRNAs, first by analysing already available datasets (two female and one male RNAseq datasets with different library preparations), and after by sequencing paired samples (two males and two females) to control for batch effects. By comparing the expression profile of *D. pseudoobscura* and *D. melanogaster* it was evident that the sex-bias of microRNAs is largely conserved between both species. This is also the case for microRNAs that were located in an autosome which became the neo-X arm after a translocation, supporting that the relative expression between males and females does not depend on the chromosomal context. In other words, male-biased microRNAs may acquire their expression profile early during evolution, but then their expression is maintained even if the chromosome acquires a sex chromosome status. The conservation of the expression bias as well as the conservation of the chromosome in which microRNAs are located provides strong evidence against demasculinization having any impact in microRNAs.

There were two exceptional cases, when analysing the paired dataset, of microRNAs strongly male-biased in *D. pseudoobscura* but strongly female-biased in *D. melanogaster*. These microRNAs belong to the mir-310~313, whose products are maternally deposited in the egg (Ninova et al. 2014; Marco 2015). In this study, to avoid the excessive contamination from maternal/embryonic microRNAs in females (Marco 2014; Ninova et al. 2014; Marco 2015), the RNA of young virgin females were sequenced. That may explain the differences observed in these two particular clusters. In any case, both microRNAs clusters are located in the homologous autosome in both species, so the observed difference is surely not a consequence of a different chromosomal context.

In the lineage of the species *Drosophila willistoni* there was an independent fusion of the Muller element D (chr3L in *D. melanogaster*) with the X chromosome, generating a neo-X similar to what happened in the pseudoobscura lineage. Of the male-biased *D. melanogaster* microRNAs, only two were located in the 3L chromosome: mir-274 and mir-276a. BLAST searches revealed that both microRNAs are conserved in *D. willistoni* as well as the neighboring region, suggesting that the synteny is conserved. However, as far as I know, these scaffolds are not mapped into the Muller elements and a detailed analysis of any potential movement out of the *D. willistoni* neo-X is not possible at the moment.

The demasculinization model described in the Introduction assumes that male-expressed (male-beneficial) genes are more likely to be evolutionarily lost if they are located in the X chromosome, as damaging mutations are often lethal in hemizygosity (Figure 6A). Hence, gene copies in the autosome are favored while the ancestral X located copy is eventually lost (Sturgill et al. 2007). Some groups have suggested that novel microRNAs are under positive selection upon emergence (Lyu et al. 2014), and therefore, they are more likely to be fixed in the X chromosome if they are male beneficial. However, once fixed, recessive deleterious mutations will be more damaging for males if the microRNA is in the X chromosome. As a matter of fact, a majority of novel microRNAs are lost within a relatively short evolutionary period (Lyu et al. 2014). The genes are copied by retroposition (Figure 6B), in which a reverse transcriptase makes a DNA copy of the processed transcript which is inserted (by a retroviral integrase) in the genome. The original copies of genes retroposed into the autosomes are eventually lost (Figure 6C). This mechanism is highly unlikely, if not impossible, in microRNAs since processed microRNA precursors leaving the nucleus (and therefore exposed to endogenous reverse transcriptases) are short (less than 100 nucleotides) RNA molecules with an internal hairpin structure (Bartel 2004), lacking a polyadenylation sequence that could be used as a primer by the endogenous reverse transcriptases (Wei et al. 2001). DNA-based gene relocation (that is, not involving retroposition) has also been described in X to autosome gene movement (Vibranovski et al. 2009). However, it was not detected in this analysis any microRNA X to autosome translocation, so DNA-based demasculinization has not been observed either, yet it will be interesting to explore microRNA inter-chromosomal movement at longer evolutionary distances (although the current assemblies do not allow to perform this analysis yet). Nevertheless, in future works DNA-based transposition should be taken into account as the most likely mechanism of microRNA interchromosomal movement.

**Figure 6.**
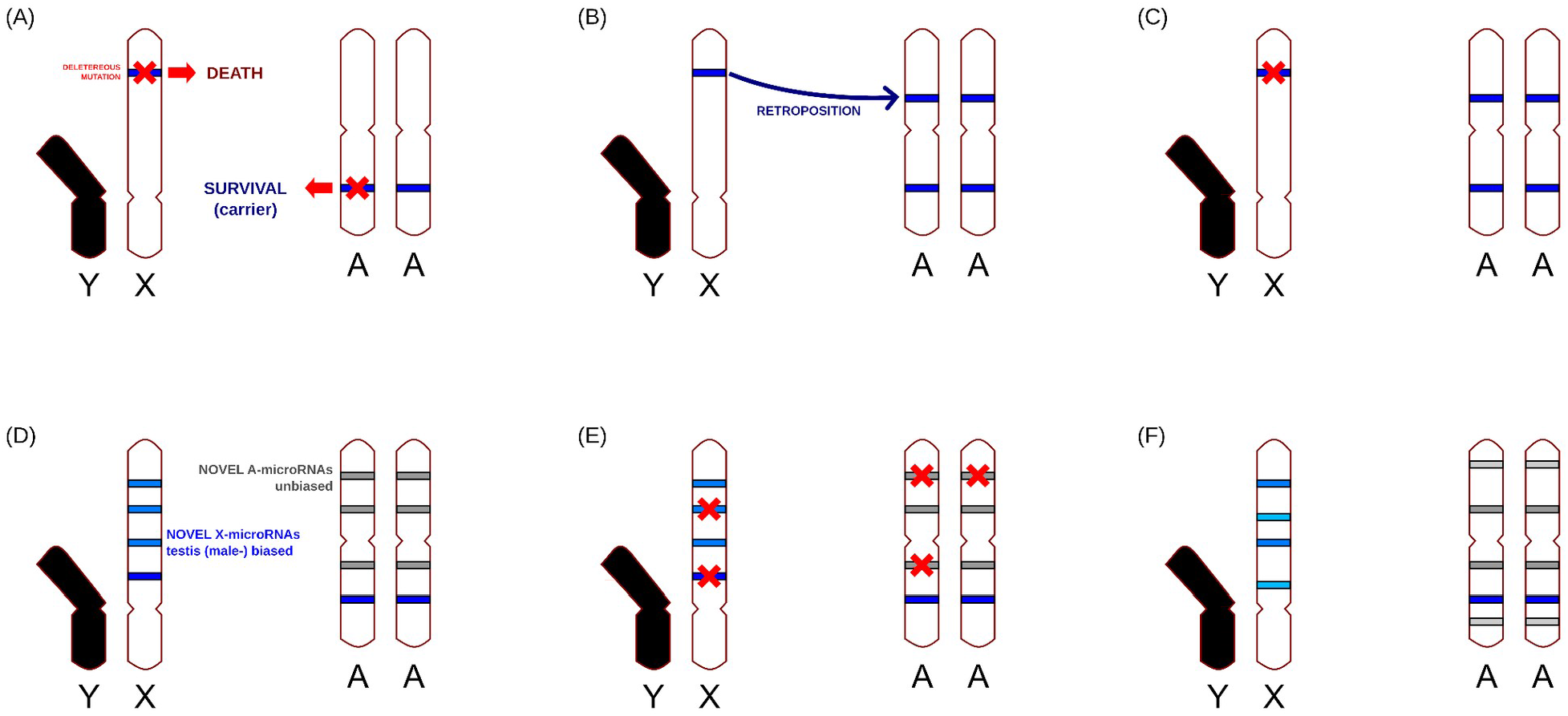
Models of protein-coding and microRNA genes evolution. (A) Effect of deleterious mutations in male genes. (B) Retroposition out of the X chromosome. (C) Deletion of the original copy of a retroposed gene. (D) Novel microRNAs emerge (light color) but in the X chromosome they tend to be highly-expressed in testis (blue). (E) Random deleterious mutations impair the function of microRNA genes, which are eventually lost. (F) As new testis-expressed microRNAs accumulate preferentially in the X chromosome, an enrichment of male-biased microRNAs in the X chromosome is observed. Details of the models are in the main text.

This work supports an alternative model of evolution. MicroRNAs have a high rate of turn-over. That is, novel microRNAs appear at high frequencies (Figure 6D), which is less common in protein-coding genes. Novel genes are often male-biased (Metta and Schlötterer 2008), and if they are in the X they are frequently highly expressed in testis (this work, see also (Mohammed et al. 2014)). Once a new microRNA transcript with male-biased expression emerges in the X chromosome, it can potentially serve as a source of novel linked microRNAs within the transcript, which may explain why evolutionarily young X-linked male-biased microRNAs tend to appear in clusters, not only in *Drosophila*, but also in mammals (Figure 5B; (Li et al. 2010; Devor et al. 2011; Marco, Ninova, Ronshaugen, et al. 2013)). This bias in gene expression of newly emerged microRNAs, together with the high rate of evolutionarily young microRNA loss (Figure 6E) eventually leads to an enrichment of male (testis) expressed microRNAs in the X chromosome (Figure 6F). Alternatively, if X-located male-biased microRNAs had an important function and, therefore, were under purifying selection, this selection is expected be significantly weaker than in protein-coding genes, as only a very small fraction of nucleotides of a whole microRNA transcript is functional (Rodriguez et al. 2004), while for protein coding genes a large fraction of exonic nucleotides code for functional amino acids. In other words, while a microRNA gene contains dozens of nucleotides necessary for the production of a mature microRNA, a protein-coding gene of the same size contains thousands of these nucleotides, and are therefore more likely to be hit by a random mutation. Either way, natural selection seems to have a limited role in the birth and maintenance of male-biased X-linked microRNAs. Future work on older neo-X chromosomes in *Drosophila* and other insects will shed further light on the validity of this model in other species and over significantly longer evolutionary periods.

It is also important to keep into consideration that male-biased expression does not necessarily mean beneficial for males, or that losing the gene males fertility will be compromised. Indeed, genes whose lack of function lead to infertility are male expressed, but the converse does not hold true, male-biased genes are not generally fertility genes (Lindsley et al. 2013). Putting all this together, this work strongly suggest that the X-chromosome is enriched in male-biased microRNAs as a consequence of mutation bias (tendency of novel microRNAs in the X to be male biased), lack of retroposition, and a conservation of the expression-bias independently of the chromosomal context. In summary, the evidence suggests that the chromosomal location of sex-biased microRNAs is non-adaptive.

## METHODS

### Sample collection

The work was done with a *Drosophila pseudoobscura* stock kindly donated by Tom Price (University of Liverpool) and Nina Wedell (University of Exeter), collected in Show Low (Arizona, USA). All flies were kept at 18°C on cornmeal based media, with 12 hours light/dark cycles. Adult males and females were collected at age 7 days. Unfertilized eggs were collected from virgin females in population cages (about 30-50 females) in cycles of 12 hours. Virgin females were allowed to lay eggs in control vials to ensure they were virgins. We discarded population cages with females coming from control vials having larvae within two weeks (in these experiments, it only happened once). Eggs were collected with a sieve and washed with wash solution (NaCl 100 mM, Triton X-100 0.7 mM) and water.

### RNA extraction and sequencing

Total RNA from four samples (two males and two females) samples was extracted using TRIzol (Life Technologies) as recommended by the manufacturer. RNA was dissolved in RNase-free water. For small RNA sequencing I used the TruSeq Small RNA Sample Preparation Kit (Illumina) to generate the cDNA library using selected constructs of sizes 145 to 160 bp in a 6% PAGE gel, and precipitated in ethanol. DNA integrity was checked with TapeStation (Agilent). Samples were sequenced in-house with an Illumina MiSeq sequencing machine.

### MicroRNA expression analysis

Reads were mapped to the *Drosophila pseudoobscura* genome sequence assembly 104 (Liao et al. 2021) with HISAT2 version 2.2.1 (Kim et al. 2019) with default parameters. Raw reads were deposited in the Gene Expression Omnibus (GEO) at NCBI with accession number GSE179989. Additionally, reads from other *D. pseudoobscura* samples were retrieved from GEO: GSE98013 (Mohammed et al. 2018) (two males and one female samples) and GSE48017 (Lyu et al. 2014) (one ovary and one testis samples) and also mapped to the same reference genome. Reads from *D. melanogaster* SRR016854, SRR018039, SRR069836 and SRR069837 (Chung et al. 2008; Roy et al. 2010) were mapped with the same method to the reference genome dm6 (Hoskins et al. 2015). Adapters were trimmed with Cutadapt 1.18 (Martin 2011) prior to mapping. Read counts were obtained with featureCounts 2.0.2 (Liao et al. 2014) using the annotation coordinates from miRBase release 21 (Kozomara et al. 2018) and the additional microRNAs originally annotated by (Mohammed et al. 2018), filtering out those mapping with 100% identity to multiple regions in the current genome assembly. MicroRNAs labeled as erroneously annotated by this study were also removed from the analysis. Differential gene expression (DGE) analyses of our *D. pseudoobscura* samples, and the *D. melanogaster* male and female available datasets, were conducted with DESeq2 version 1.24.0 (Love et al. 2014: 2) with local fitting. Alternatively, DGE was also done with limma version 4.2.0 (Ritchie et al. 2015) using TMM normalization and voom transformation (Law et al. 2014). For both DGE methods the model used was expression ~ batch + sex, where batch is the sequencing run. In the fold-change comparison between chromosomes and novel and conserved microRNAs, only those with at least one read count in one of the samples in each sex and a base mean (from DESeq2) of at least one were considered.

### Homology analysis

Annotated microRNAs from both species were compared via BLAST 2.9.0 (Altschul et al. 1997) using the R wrapper rBLAST 0.992 (https://github.com/mhahsler/rBLAST/): blastn with word size 10 and E-value threshold of 0.1. Pairs of similar sequences between species were built into a graph with igraph 1.2.11 (https://igraph.org) and connected graphs were extracted as similarity groups (potentially homology groups). For each similarity group a sequence alignment with MAFFT 2.9.0 (Katoh and Toh 2008) and a neighbor-joining tree (Saitou and Nei 1987) using uncorrected distances with ips 0.0.11 (https://www.rdocumentation.org/packages/ips) were built for visual inspection. To identify potential, and previously missed, microRNAs, *D. melanogaster* microRNA precursors were compared against the *D. pseudoobscura* genome with blastn (word size 10, Evalue threshold of 0.01). Alignments of precursor microRNAs of length 60 or over were further considered. Lineage specific (novel) microRNAs were those described by (Mohammed et al. 2018).

All data generated and scripts to reproduce the full analysis is available from GitHub: https://github.com/antoniomarco/miRpseudoobscura

## ACKNOWLEDGEMENTS

I would like to thank Tom Price and Nina Wedell for sharing their *Drosophila pseudoobscura* stocks. I am also grateful to J. J. Emerson for helping me to understand the chromosome assembly of the most recent *D. pseudoobscura* genome assembly. This work was supported by the University of Essex.

