## Supplementary File 1 for "The chromosomal distribution of sex-biased microRNAs in *Drosophila* is non-adaptive"

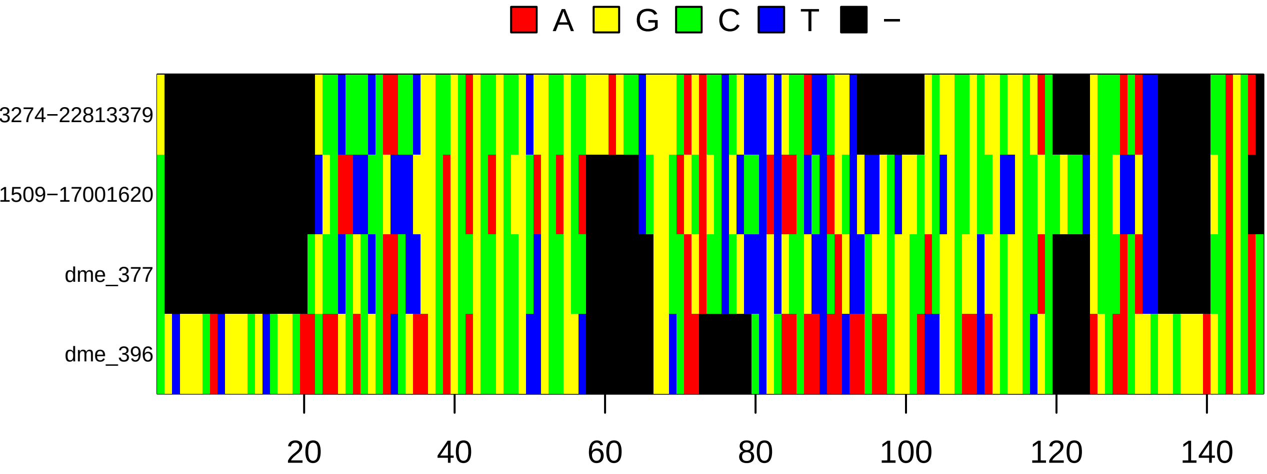
A
G
C
T
−
3274−22813379
1509−17001620
dme_377
dme_396
20
40
60
80
100
120
140

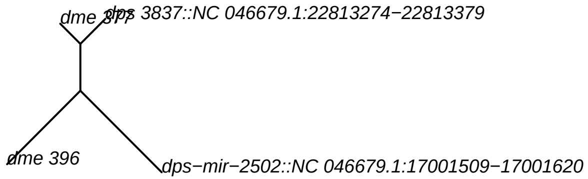
dme 377
dps 3837::NC 046679.1:22813274−22813379
dme 396
dps−mir−2502::NC 046679.1:17001509−17001620

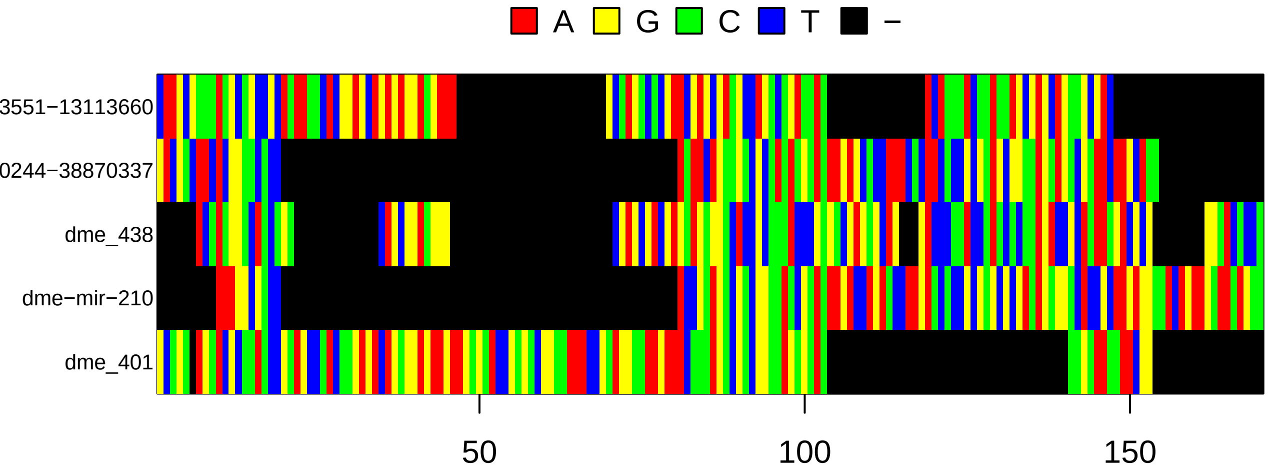
A
G
C
T
−
3551−13113660
0244−38870337
dme_438
dme−mir−210
dme_401
50
100
150

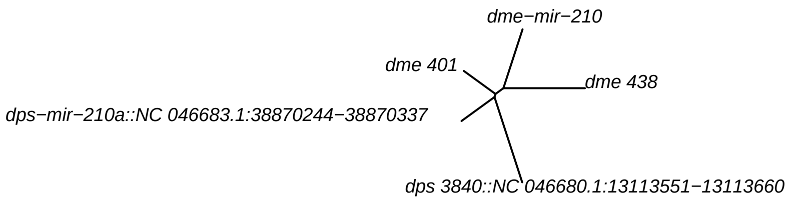
dme−mir−210
dme 401
dme 438
dps−mir−210a::NC 046683.1:38870244−38870337
dps 3840::NC 046680.1:13113551−13113660

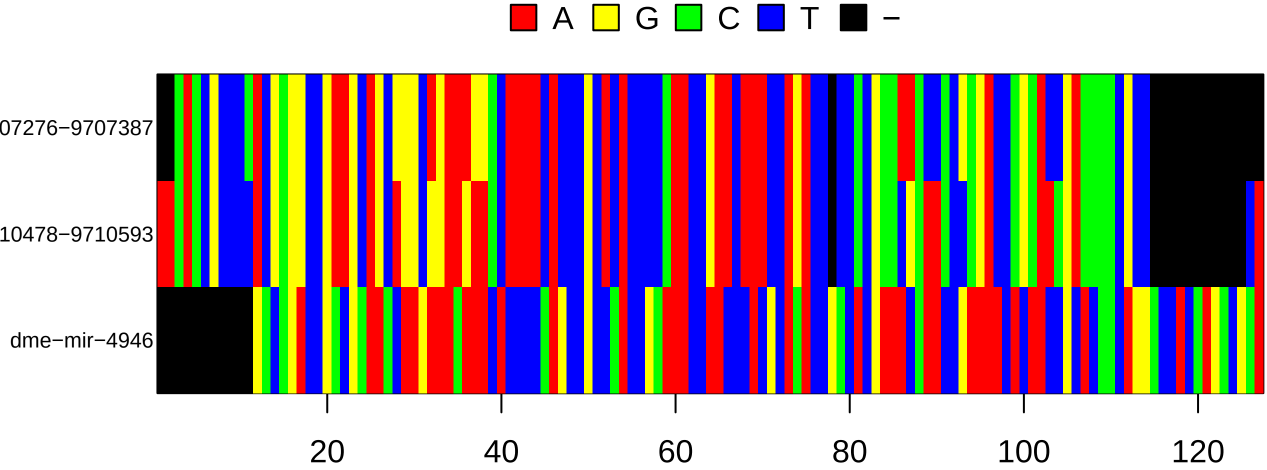
A
G
C
T
−
07276−9707387
10478−9710593
dme−mir−4946
20
40
60
80
100
120

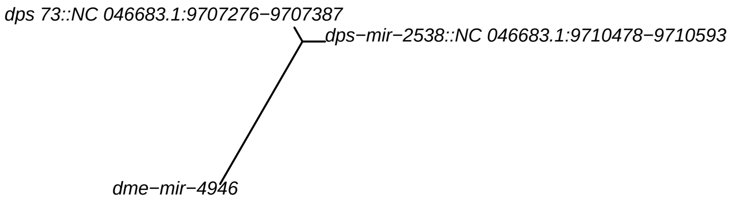
dps 73::NC 046683.1:9707276−9707387
dps−mir−2538::NC 046683.1:9710478−9710593
dme−mir−4946

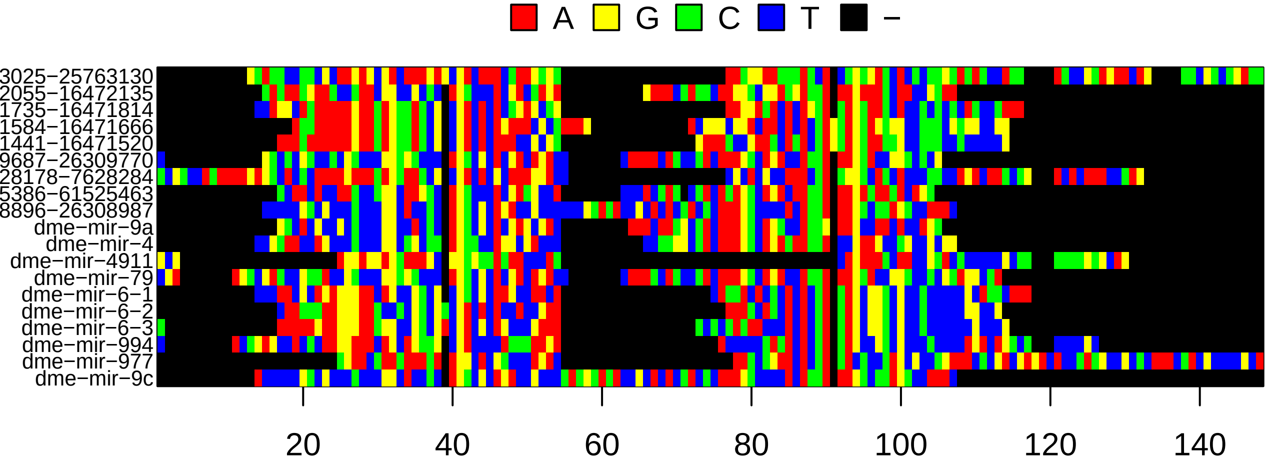
A
G
C
T
−
3025−25763130
2055−16472135
1735−16471814
1584−16471666
1441−16471520
9687−26309770
28178−7628284
5386−61525463
8896−26308987
dme−mir−9a
dme−mir−4
dme−mir−4911
dme−mir−79
dme−mir−6−1
dme−mir−6−2
dme−mir−6−3
dme−mir−994
dme−mir−977
dme−mir−9c
20
40
60
80
100
120
140

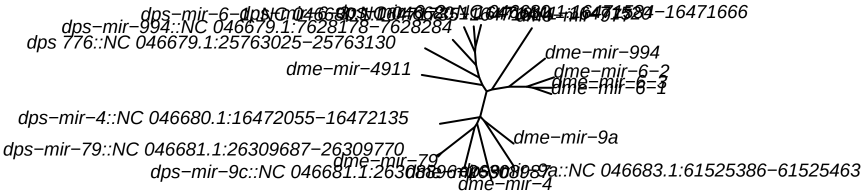
dps−mir−994::NC 046679.1:7628178−7628284
dps 776::NC 046679.1:25763025−25763130
dme−mir−4911
dps−mir−4::NC 046680.1:16472055−16472135
dps−mir−79::NC 046681.1:26309687−26309770
dme−mir−994
dme−mir−6−2
dme−mir−6−1
dme−mir−9a
dme−mir−79
dme−mir−4

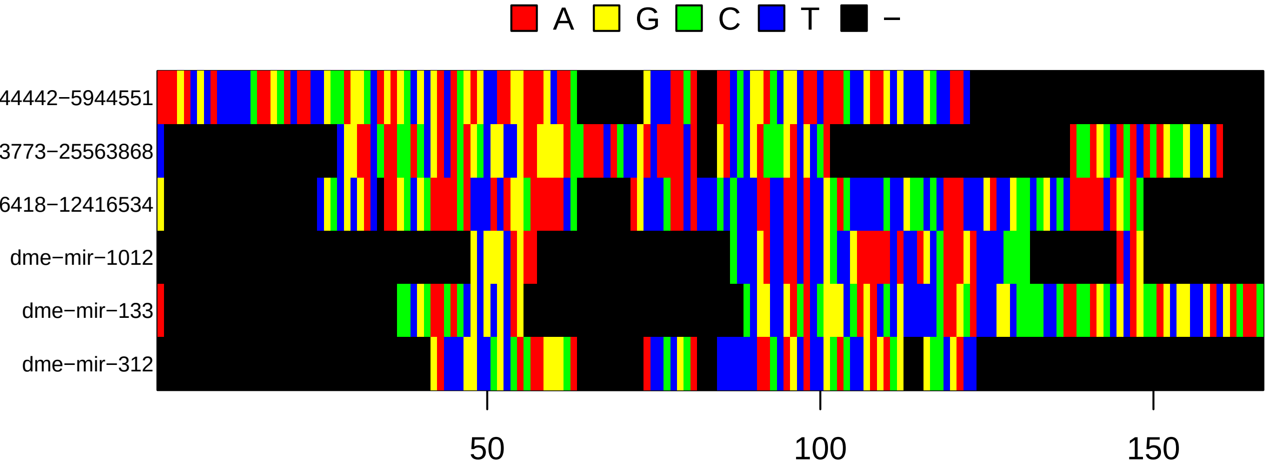

A
G
C
T
−
44442−5944551
3773−25563868
6418−12416534
dme−mir−1012
dme−mir−133
dme−mir−312
50
100
150


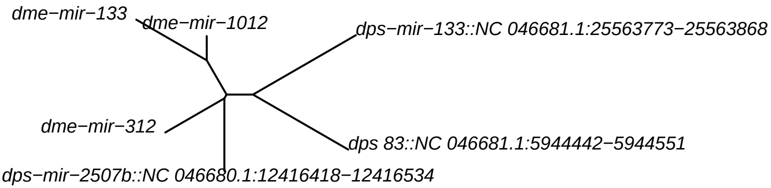

dme−mir−133
dme−mir−1012
dps−mir−133::NC 046681.1:25563773−25563868
dme−mir−312
dps 83::NC 046681.1:5944442−5944551
dps−mir−2507b::NC 046680.1:12416418−12416534

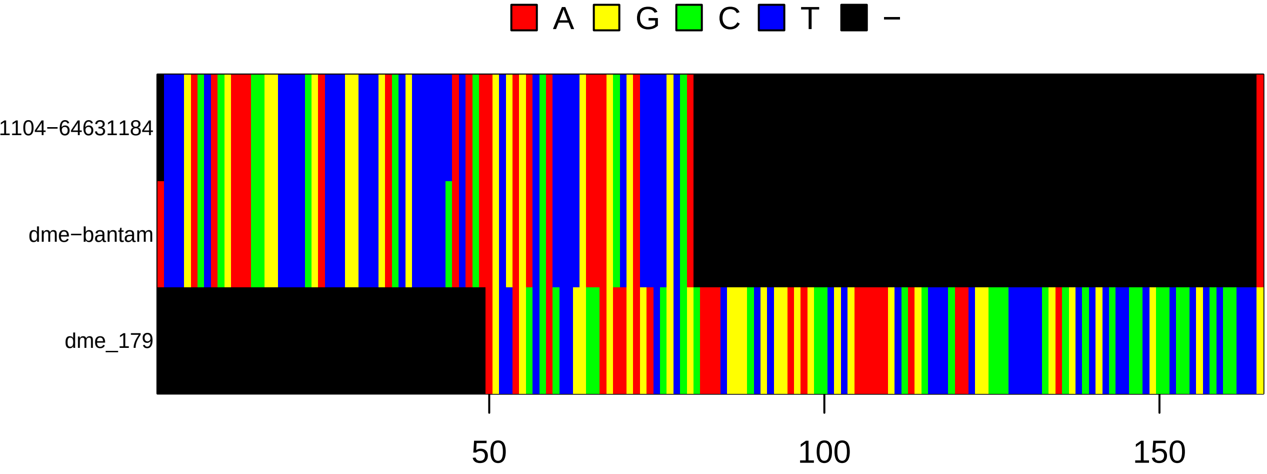
A
G
C
T
−
1104−64631184
dme−bantam
dme_179
50
100
150

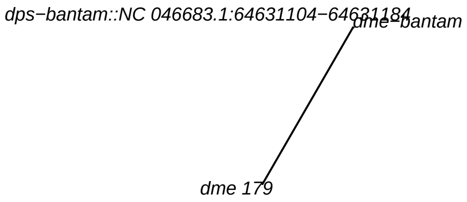
dps−bantam::NC 046683.1:64631104−64631184
dme−bantam
dme 179

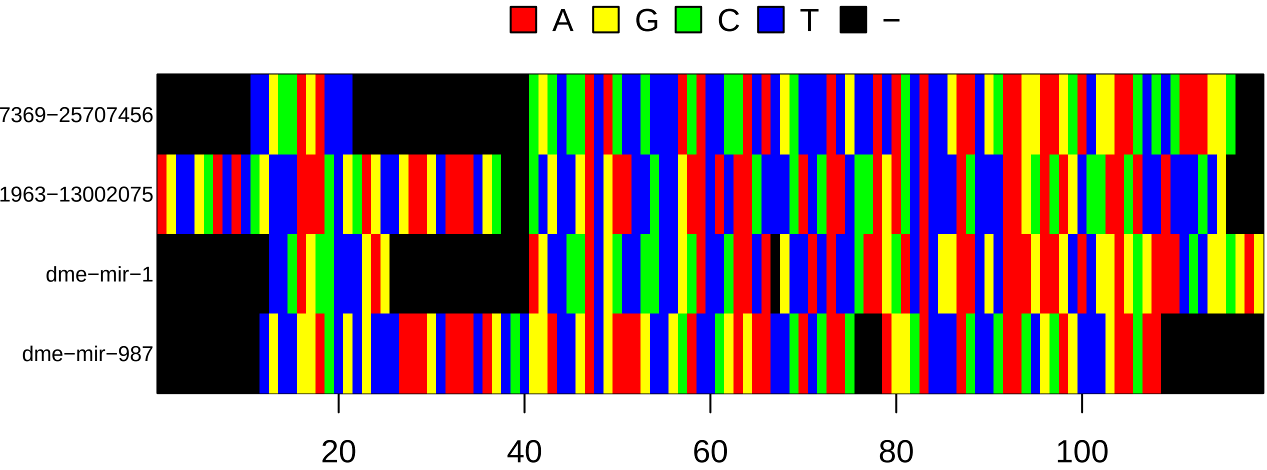

A
G
C
T
−
7369−25707456
1963−13002075
dme−mir−1
dme−mir−987
20
40
60
80
100


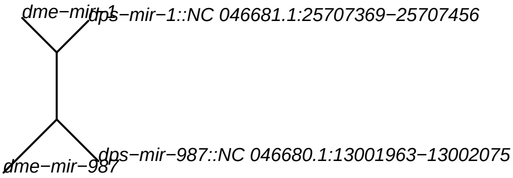

dme−mir−1
dps−mir−1::NC 046681.1:25707369−25707456
dme−mir−987
dps−mir−987::NC 046680.1:13001963−13002075

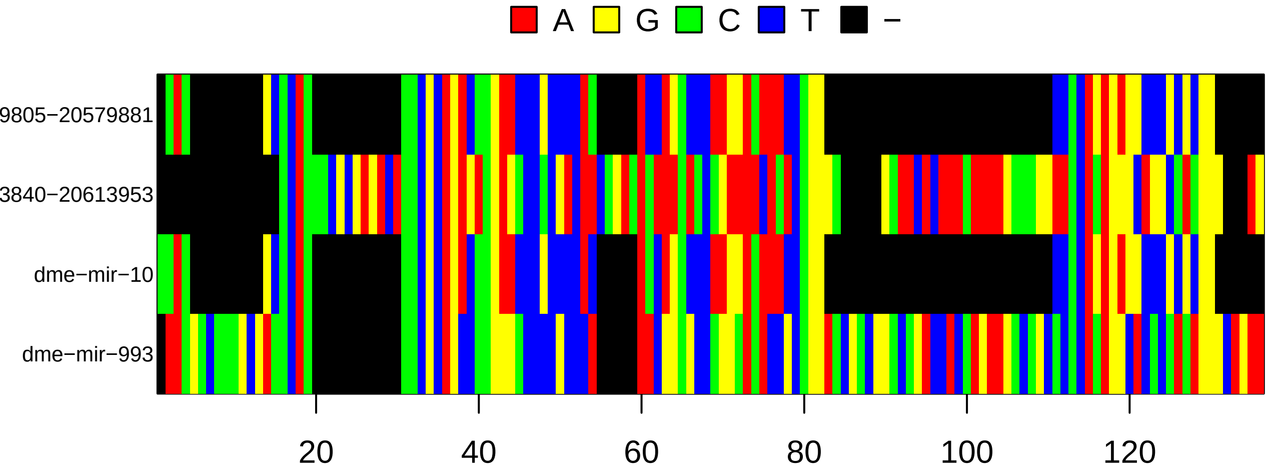
A
G
C
T
−
9805−20579881
3840−20613953
dme−mir−10
dme−mir−993
20
40
60
80
100
120

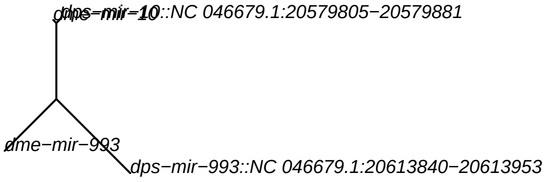
dme−mir−10
dps−mir−10::NC 046679.1:20579805−20579881
dme−mir−993
dps−mir−993::NC 046679.1:20613840−20613953

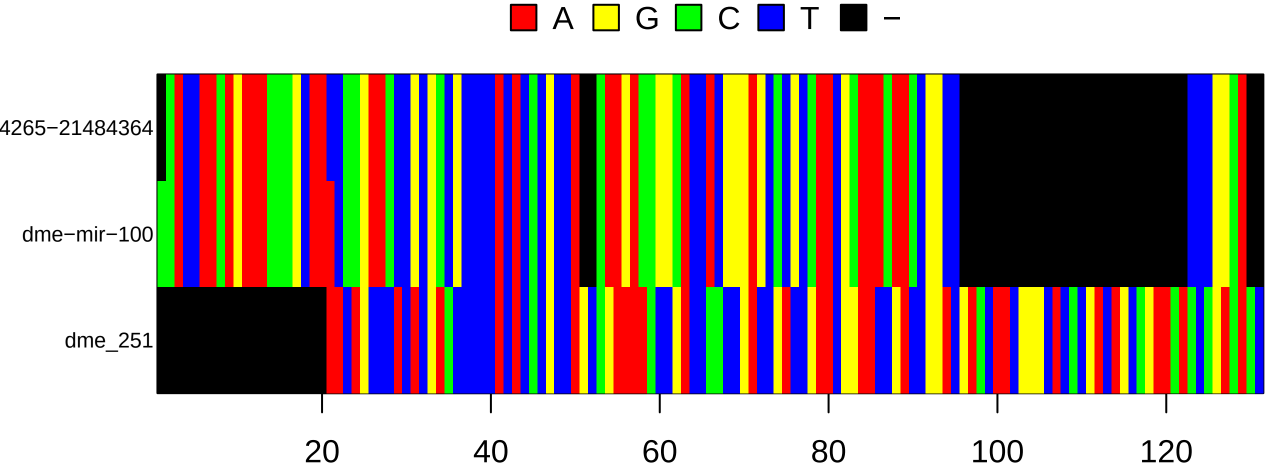
A
G
C
T
−
4265−21484364
dme−mir−100
dme_251
20
40
60
80
100
120

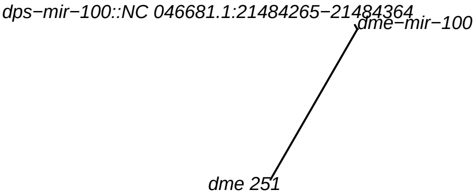
dps−mir−100::NC 046681.1:21484265−21484364
dme−mir−100
dme 251

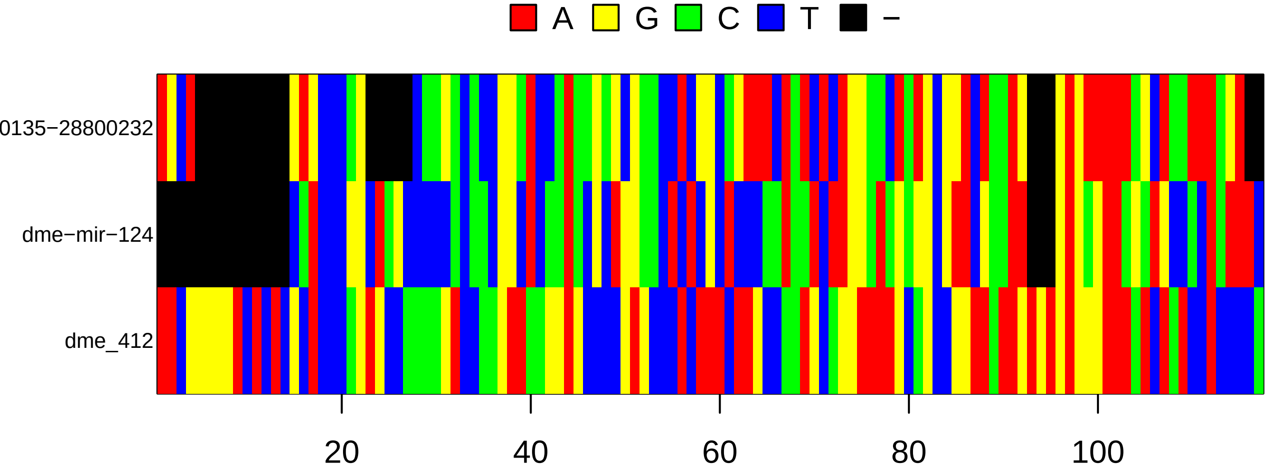
A
G
C
T
−
0135−28800232
dme−mir−124
dme_412
20
40
60
80
100

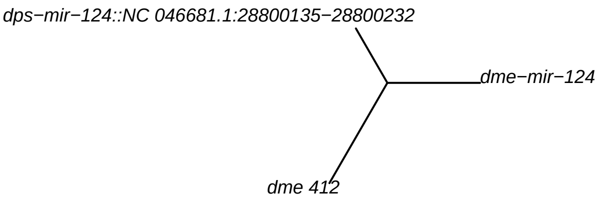
dps−mir−124::NC 046681.1:28800135−28800232
dme−mir−124
dme 412

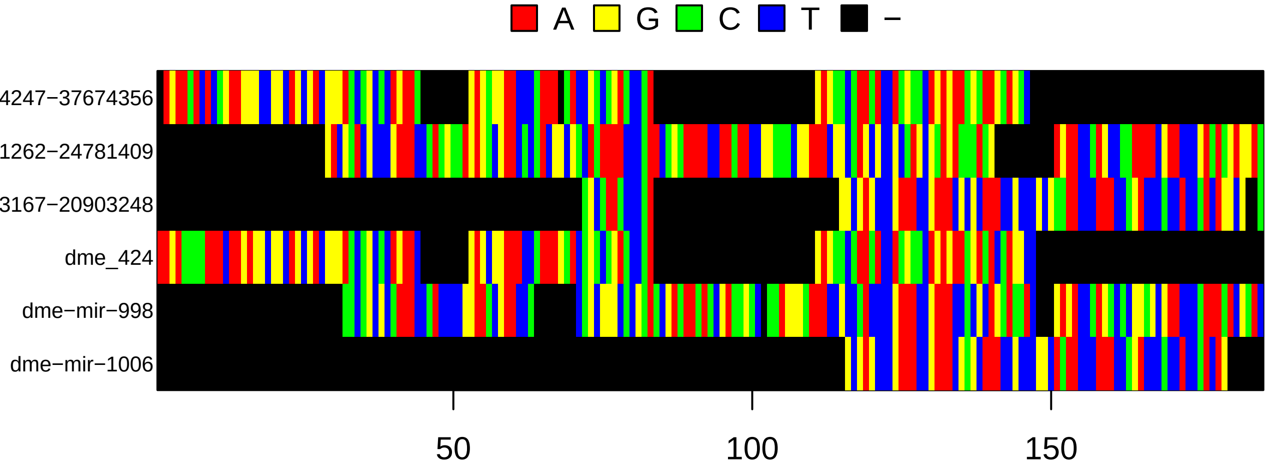

A
G
C
T
−
4247−37674356
1262−24781409
3167−20903248
dme_424
dme−mir−998
dme−mir−1006
50
100
150


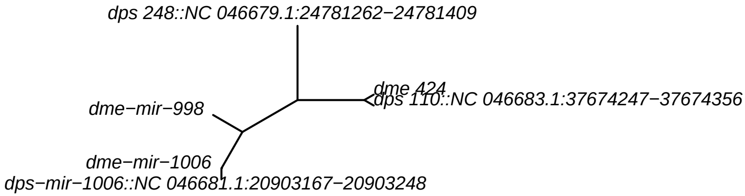

dps 248::NC 046679.1:24781262−24781409
dme 424
dps 110::NC 046683.1:37674247−37674356
dme−mir−998
dme−mir−1006
dps−mir−1006::NC 046681.1:20903167−20903248

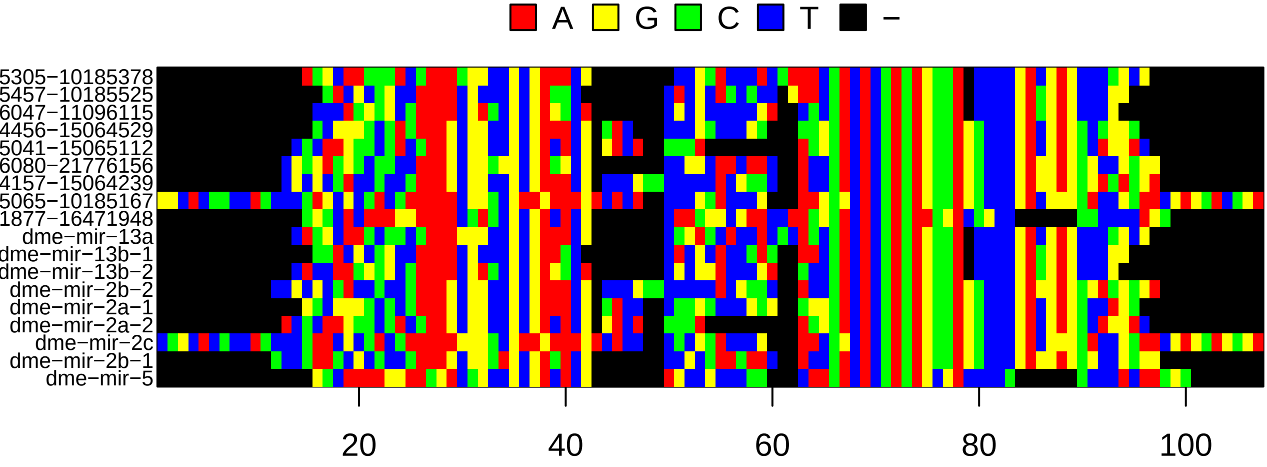
A
G
C
T
–
5305−10185378
5457−10185525
6047−11096115
4456−15064529
5041−15065112
6080−21776156
4157−15064239
5065−10185167
1877−16471948
dme−mir−13a
dme−mir−13b−1
dme−mir−13b−2
dme−mir−2b−2
dme−mir−2a−1
dme−mir−2a−2
dme−mir−2c
dme−mir−2b−1
dme−mir−5
20
40
60
80
100

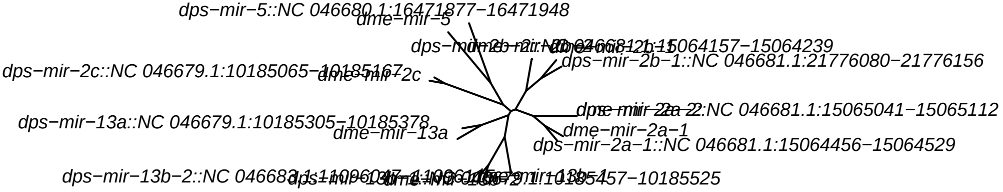
dps−mir−5::NC 046680.1:16471877−16471948
dme−mir−5
dps−mir−2b−1::NC 046681.1:21776080−21776156
dps−mir−2c::NC 046679.1:10185065−10185167
dme−mir−2c
dps−mir−13a::NC 046679.1:10185305−10185378
dme−mir−13a
dme−mir−2a−1
dps−mir−2a−1::NC 046681.1:15064456−15064529

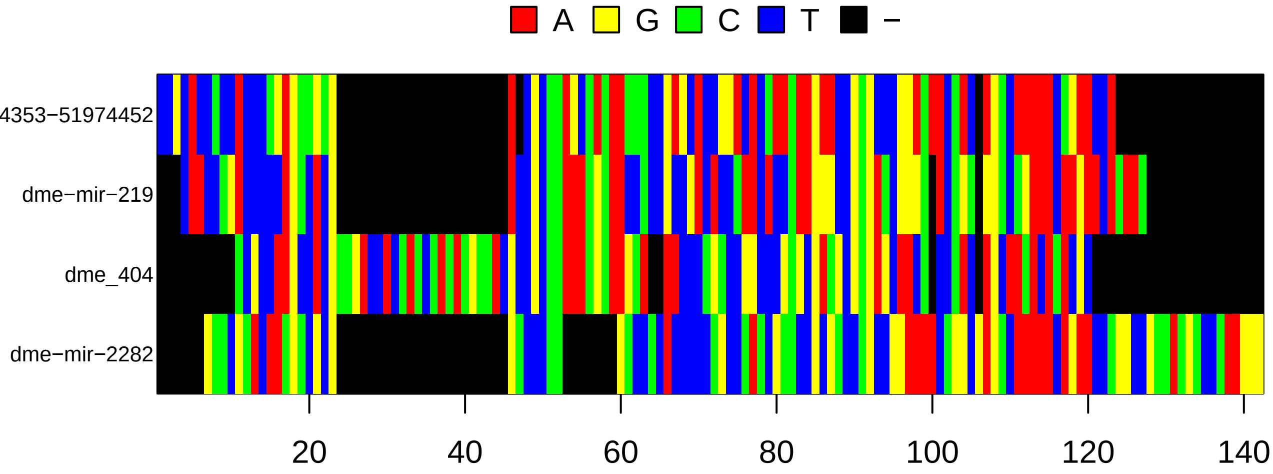
A
G
C
T
−
4353−51974452
dme−mir−219
dme_404
dme−mir−2282
20
40
60
80
100
120
140

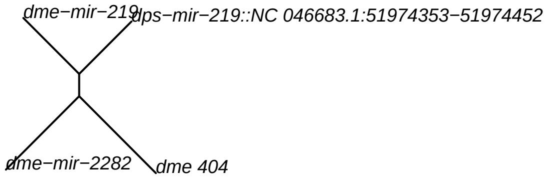
dme−mir−219
dps−mir−219::NC 046683.1:51974353−51974452
dme−mir−2282
dme 404

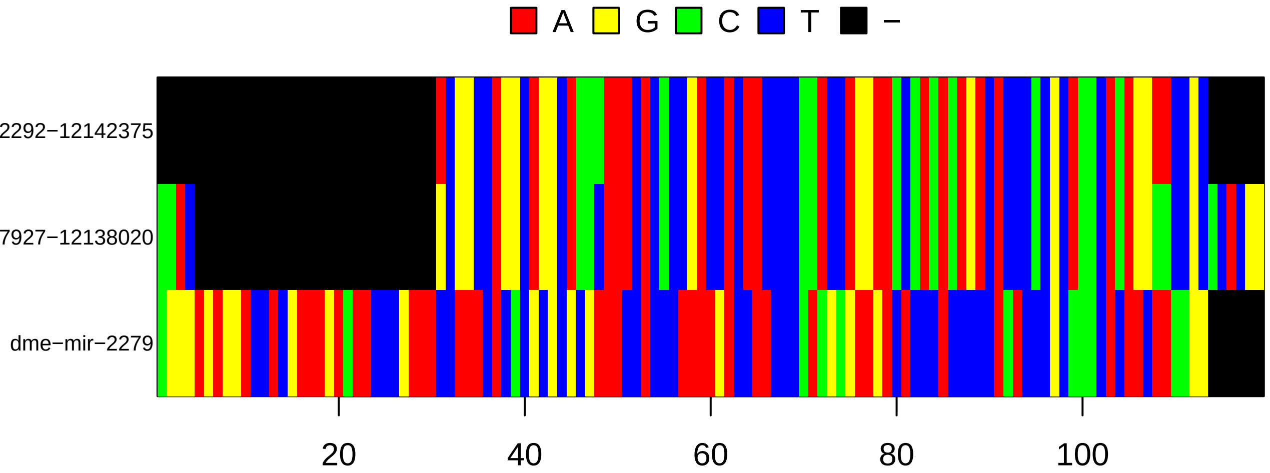
A
G
C
T
−
2292−12142375
7927−12138020
dme−mir−2279
20
40
60
80
100

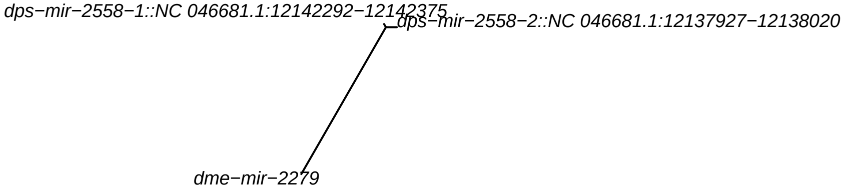
dps−mir−2558−1::NC 046681.1:12142292−12142375
dps−mir−2558−2::NC 046681.1:12137927−12138020
dme−mir−2279

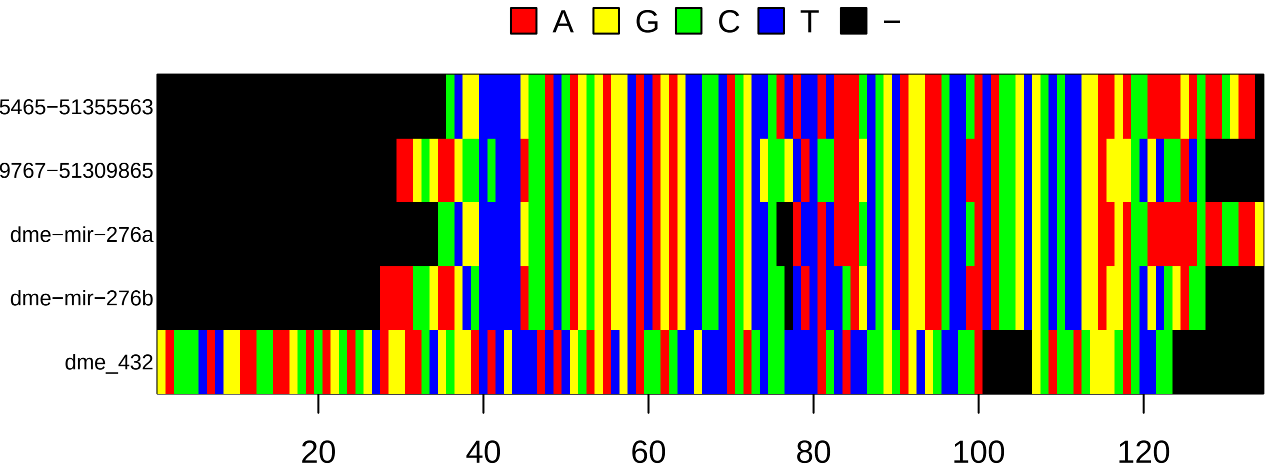
A
G
C
T
−
5465−51355563
9767−51309865
dme−mir−276a
dme−mir−276b
dme_432
20
40
60
80
100
120

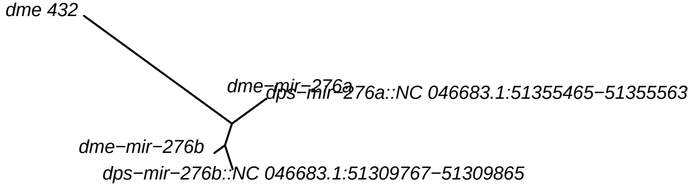
dme 432
dme−mir−276a
dps−mir−276a::NC 046683.1:51355465−51355563
dme−mir−276b
dps−mir−276b::NC 046683.1:51309767−51309865

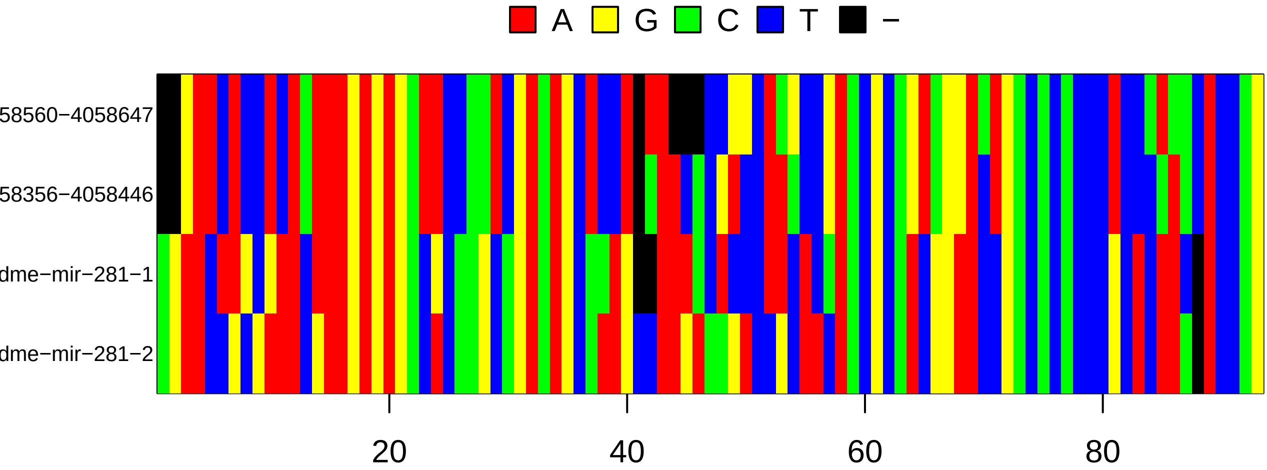
A
G
C
T
−
58560−4058647
58356−4058446
dme−mir−281−1
dme−mir−281−2
20
40
60
80

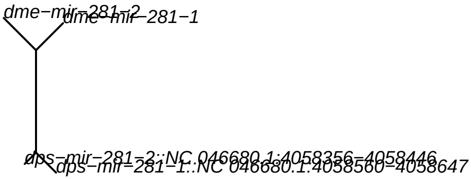
dme−mir−281−2
dme−mir−281−1
dps−mir−281−2::NC 046680.1:4058356−4058446
dps−mir−281−1::NC 046680.1:4058560−4058647

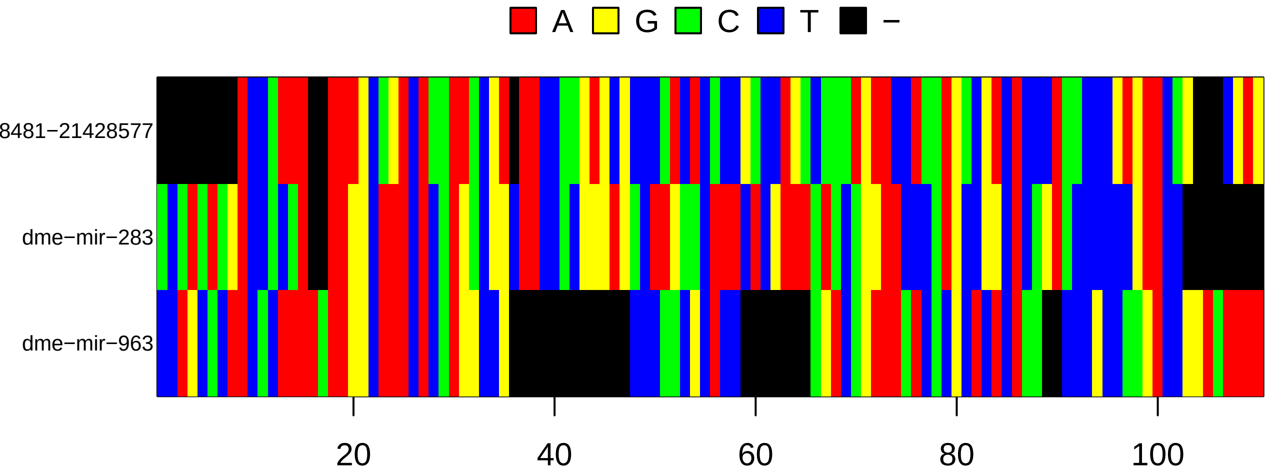
A
G
C
T
−
8481−21428577
dme−mir−283
dme−mir−963
20
40
60
80
100

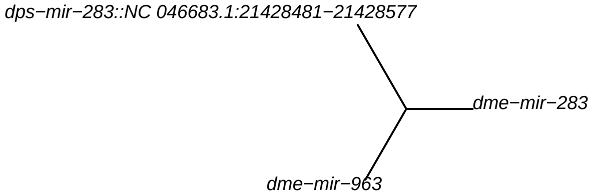
dps−mir−283::NC 046683.1:21428481−21428577
dme−mir−283
dme−mir−963

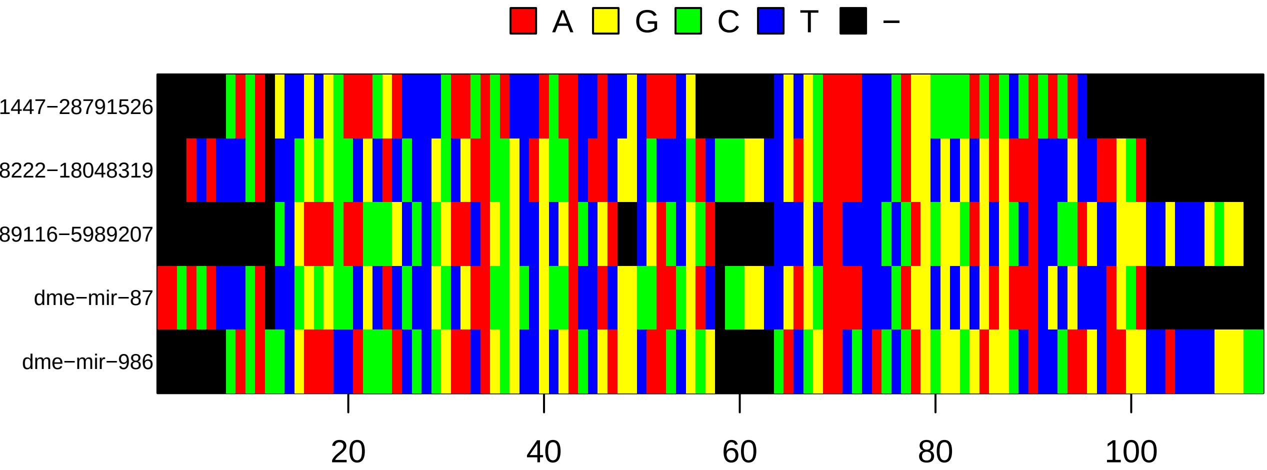

A
G
C
T
−
1447−28791526
8222−18048319
89116−5989207
dme−mir−87
dme−mir−986
20
40
60
80
100


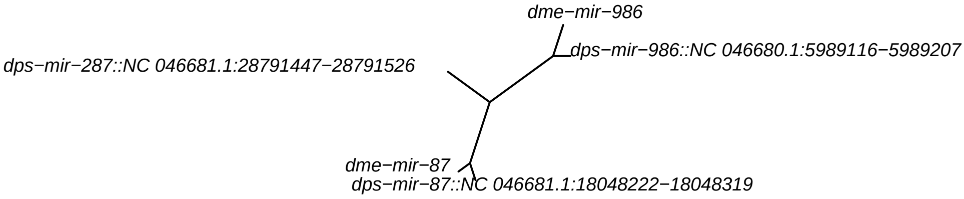

dme−mir−986
dps−mir−986::NC 046680.1:5989116−5989207
dps−mir−287::NC 046681.1:28791447−28791526
dme−mir−87
dps−mir−87::NC 046681.1:18048222−18048319

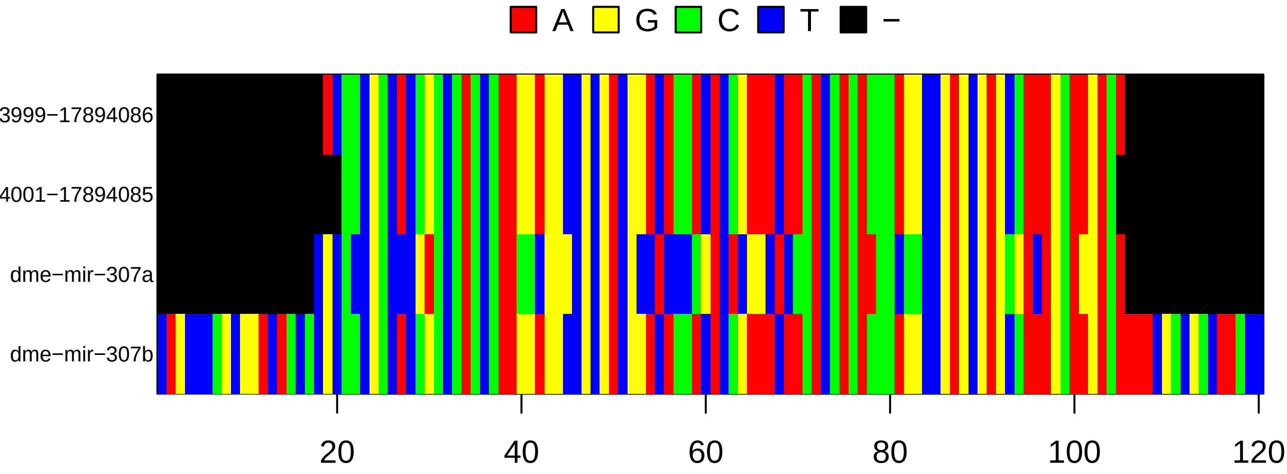
A
G
C
T
−
3999−17894086
4001−17894085
dme−mir−307a
dme−mir−307b
20
40
60
80
100
120

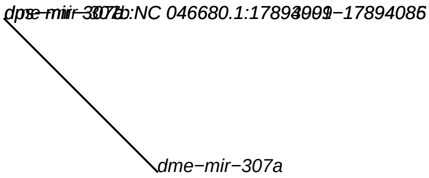
dme−mir−307a

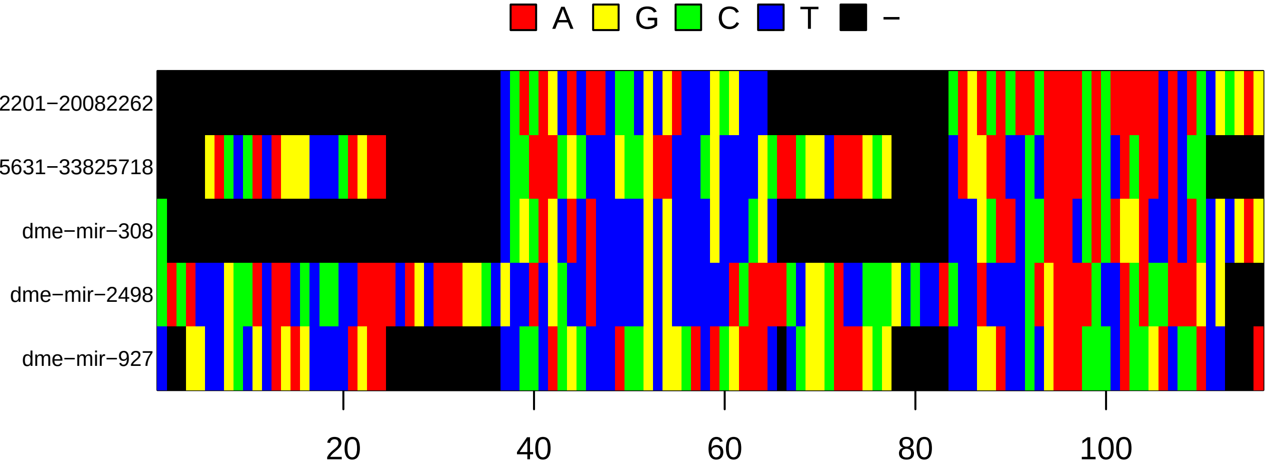
A
G
C
T
−
2201−20082262
5631−33825718
dme−mir−308
dme−mir−2498
dme−mir−927
20
40
60
80
100

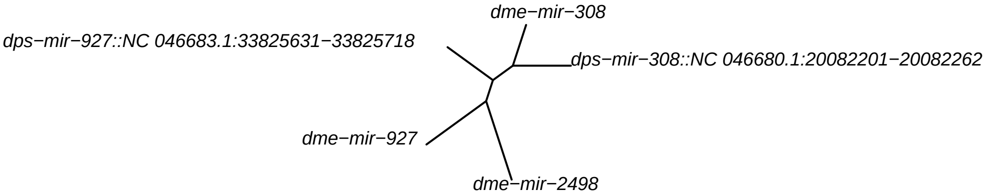
dme−mir−308
dps−mir−927::NC 046683.1:33825631−33825718
dps−mir−308::NC 046680.1:20082201−20082262
dme−mir−927
dme−mir−2498

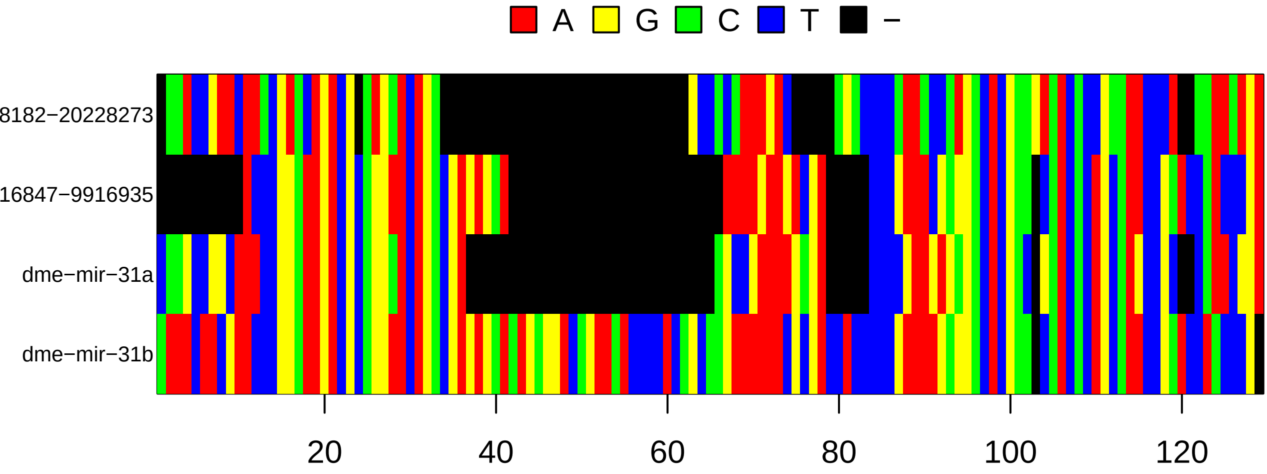
A
G
C
T
−
8182−20228273
16847−9916935
dme−mir−31a
dme−mir−31b
20
40
60
80
100
120

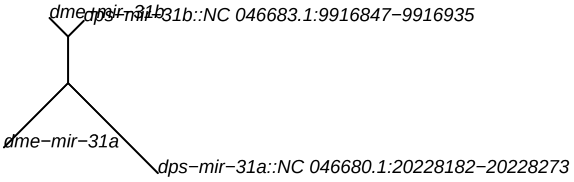
dme−mir−31b
dps−mir−31b::NC 046683.1:9916847−9916935
dme−mir−31a
dps−mir−31a::NC 046680.1:20228182−20228273

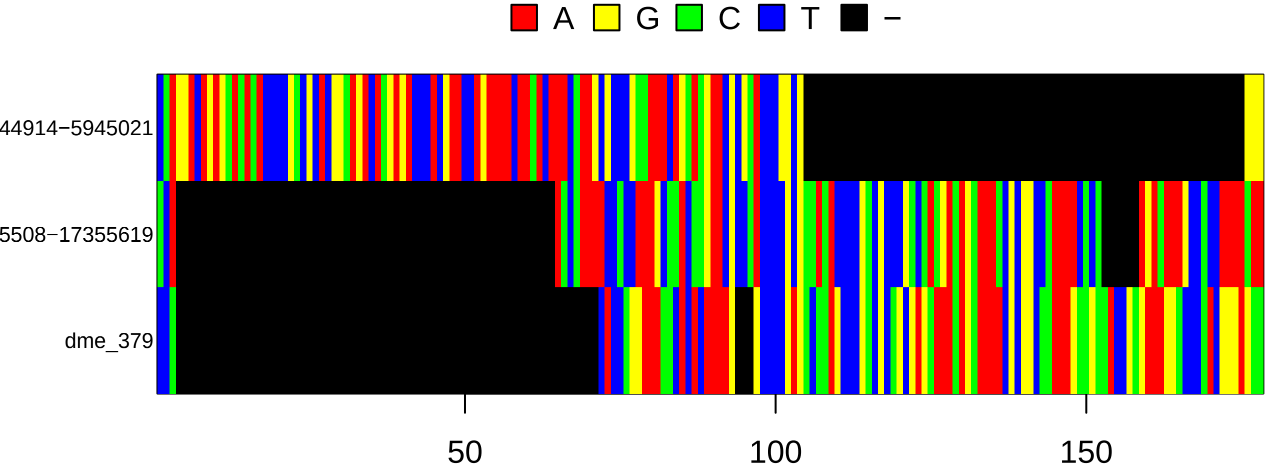
A
G
C
T
−
44914−5945021
5508−17355619
dme_379
50
100
150

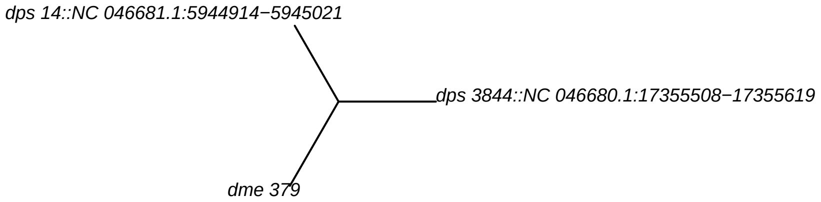
dps 14::NC 046681.1:5944914−5945021
dps 3844::NC 046680.1:17355508−17355619
dme 379

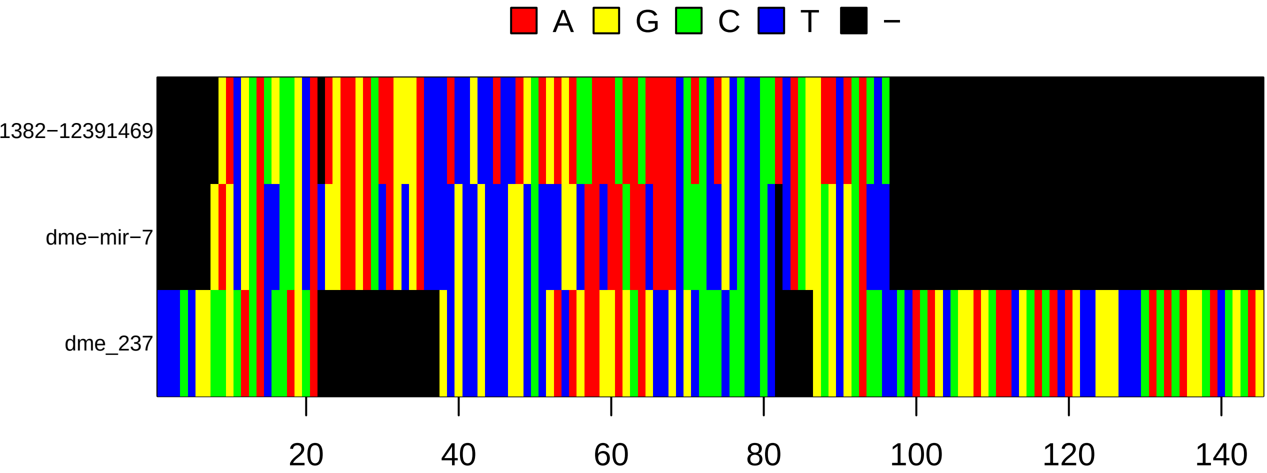

A
G
C
T
−
1382−12391469
dme−mir−7
dme_237
20
40
60
80
100
120
140


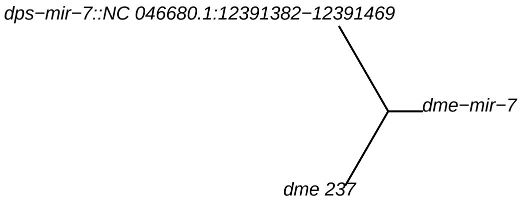

dps−mir−7::NC 046680.1:12391382−12391469
dme−mir−7
dme 237

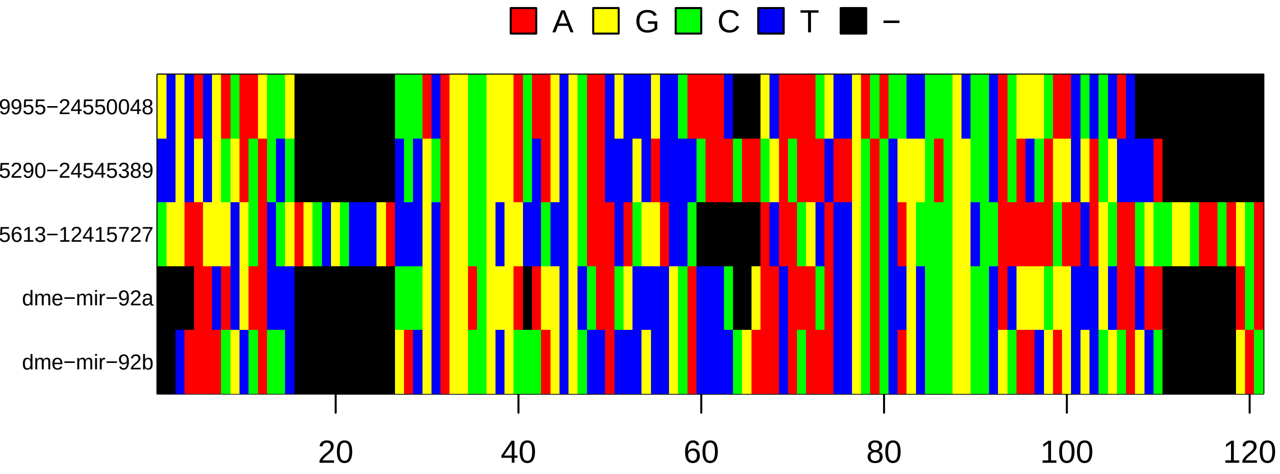

A
G
C
T
−
9955−24550048
5290−24545389
5613−12415727
dme−mir−92a
dme−mir−92b
20
40
60
80
100
120


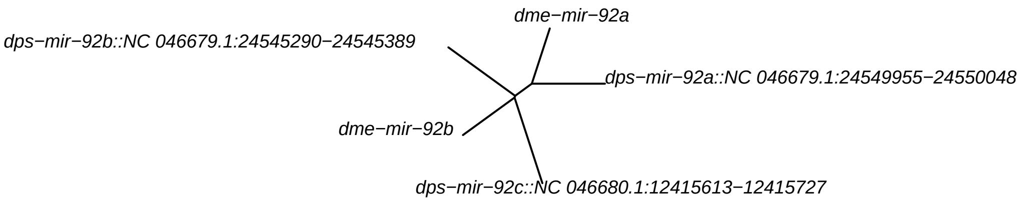

dme−mir−92a
dps−mir−92b::NC 046679.1:24545290−24545389
dps−mir−92a::NC 046679.1:24549955−24550048
dme−mir−92b
dps−mir−92c::NC 046680.1:12415613−12415727

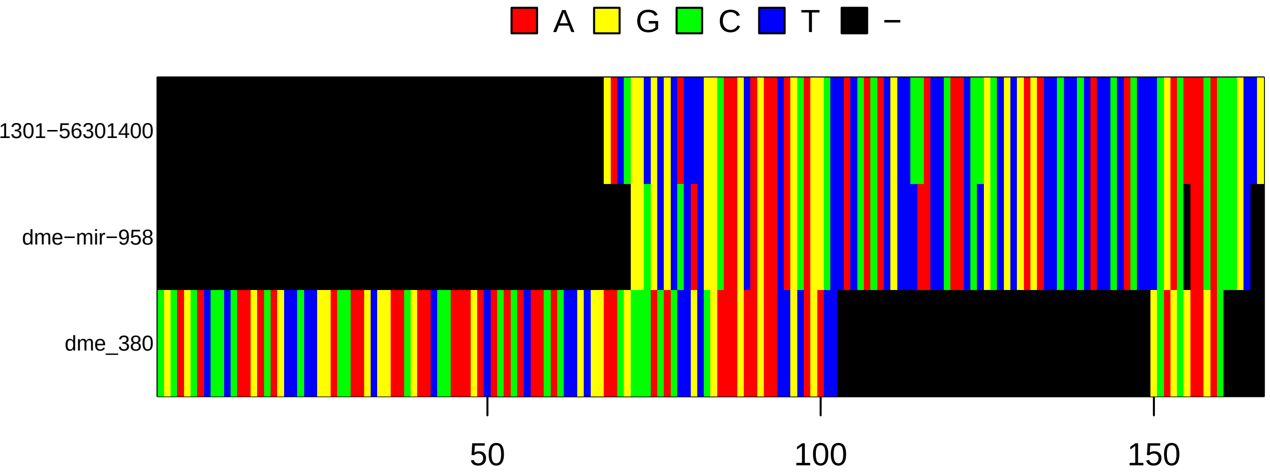
A
G
C
T
−
1301−56301400
dme−mir−958
dme_380
50
100
150

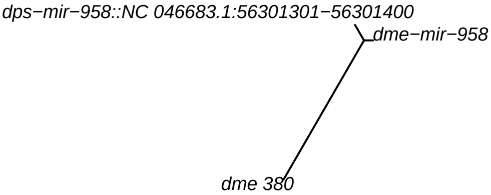
dps−mir−958::NC 046683.1:56301301−56301400
dme−mir−958
dme 380

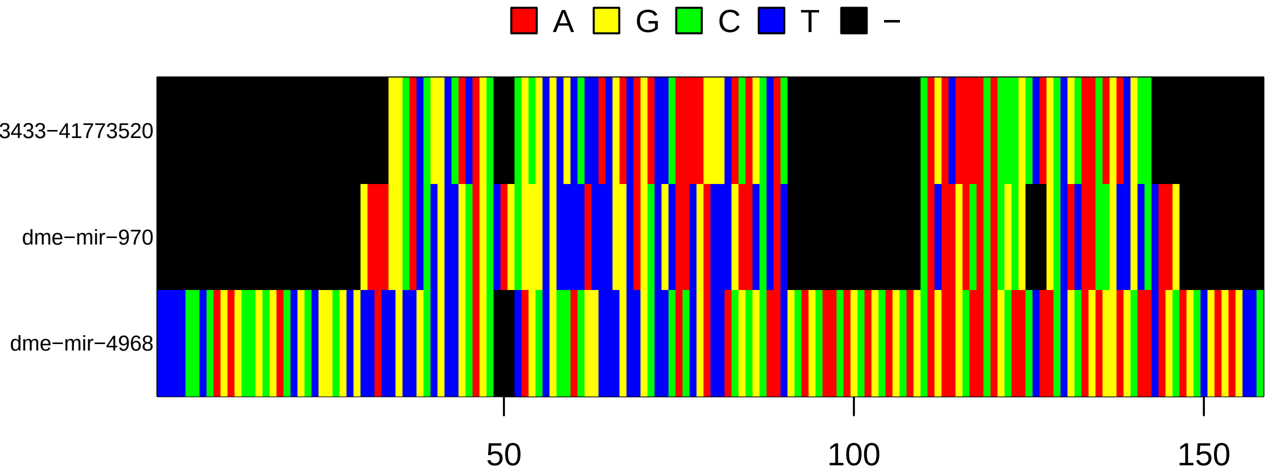
A
G
C
T
−
3433−41773520
dme−mir−970
dme−mir−4968
50
100
150

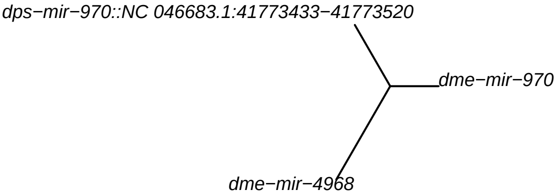
dps−mir−970::NC 046683.1:41773433−41773520
dme−mir−970
dme−mir−4968

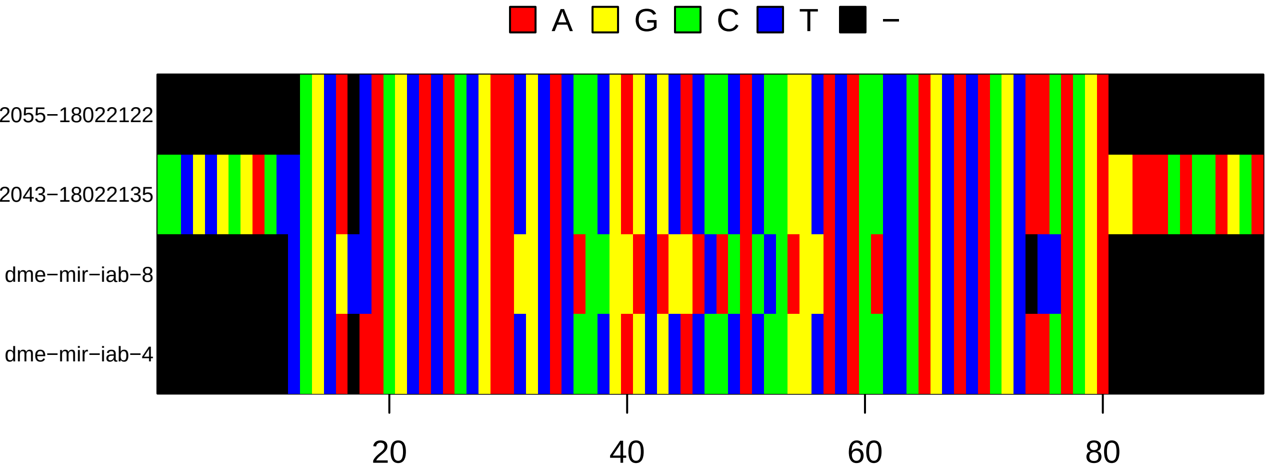
A
G
C
T
−
2055−18022122
2043−18022135
dme−mir−iab−8
dme−mir−iab−4
20
40
60
80

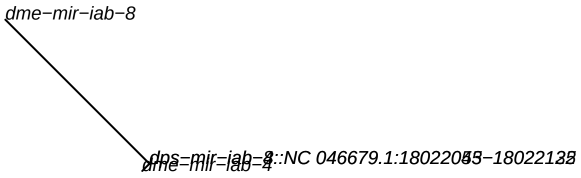
dme−mir−iab−8
dme−mir−iab−4

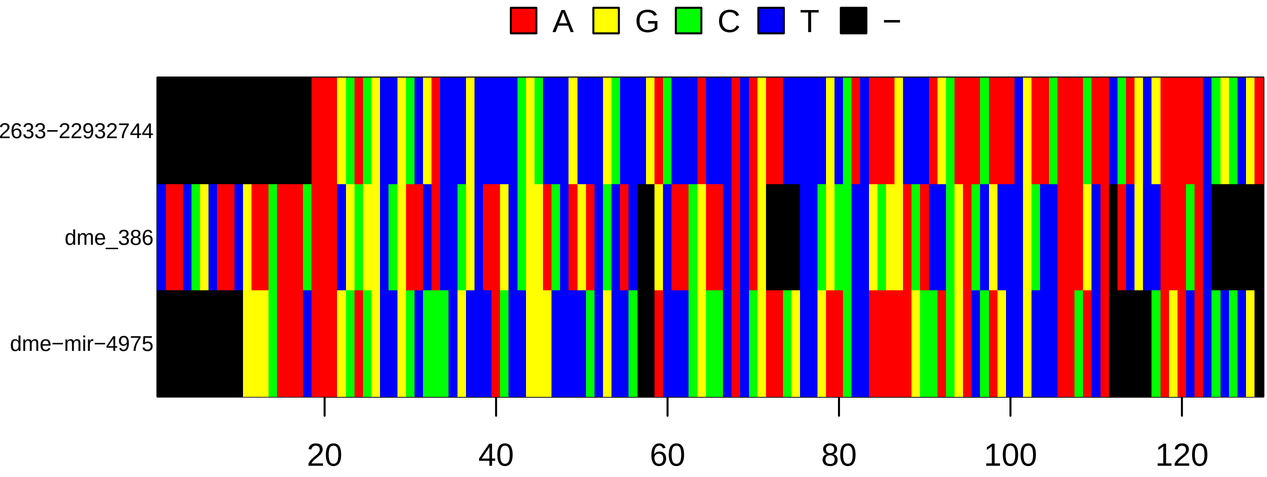
A
G
C
T
−
2633−22932744
dme_386
dme−mir−4975
20
40
60
80
100
120

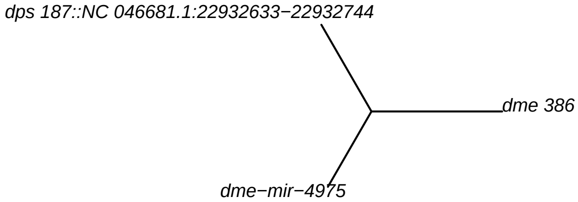
dps 187::NC 046681.1:22932633−22932744
dme 386
dme−mir−4975

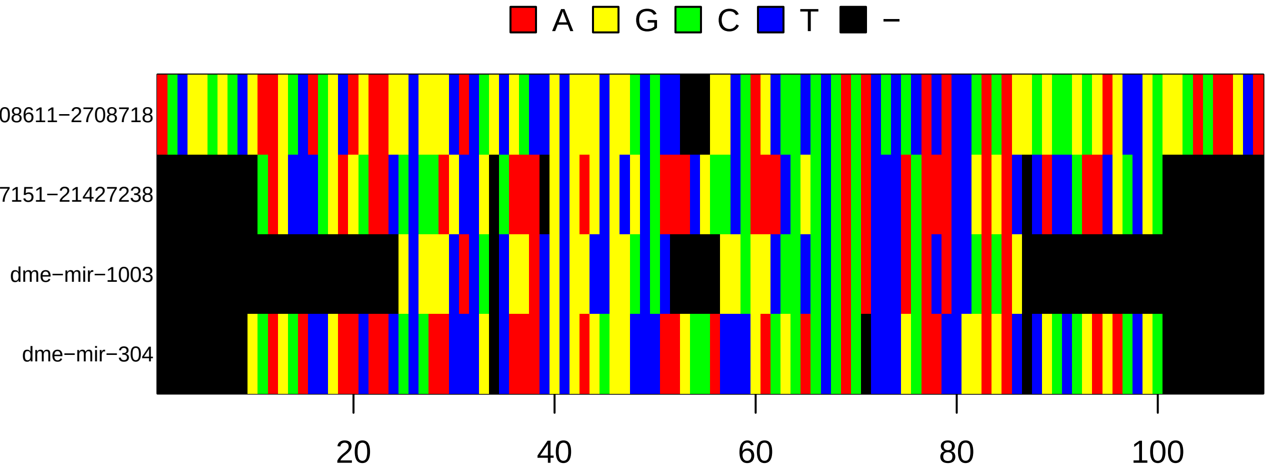
A
G
C
T
−
08611−2708718
7151−21427238
dme−mir−1003
dme−mir−304
20
40
60
80
100

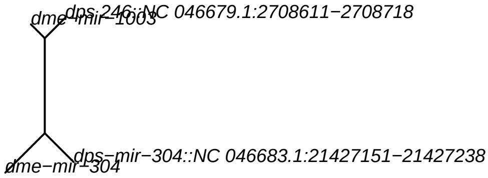
dps-246::NC 046679.1:2708611−2708718
dme−mir−1003
dps−mir−304::NC 046683.1:21427151−21427238
dme−mir−304

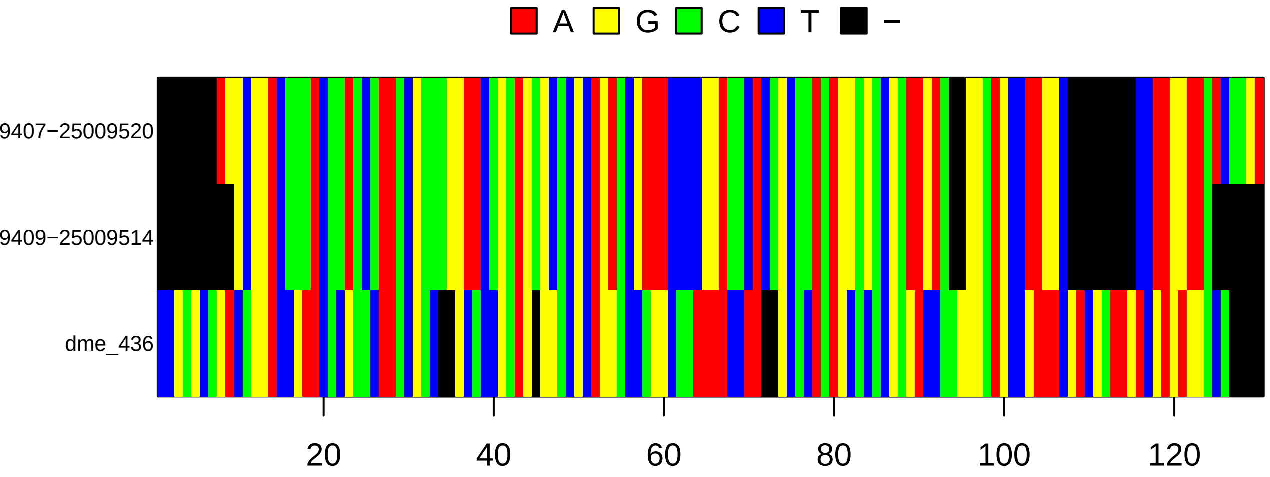
A
G
C
T
−
9407−25009520
9409−25009514
dme_436
20
40
60
80
100
120

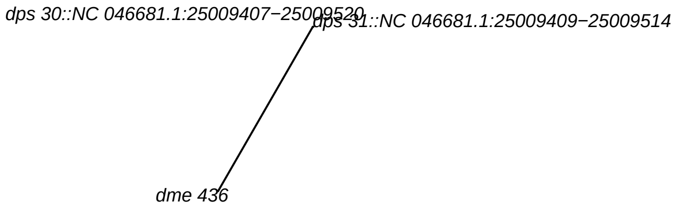
dps 30::NC 046681.1:25009407−25009520
dps 31::NC 046681.1:25009409−25009514
dme 436

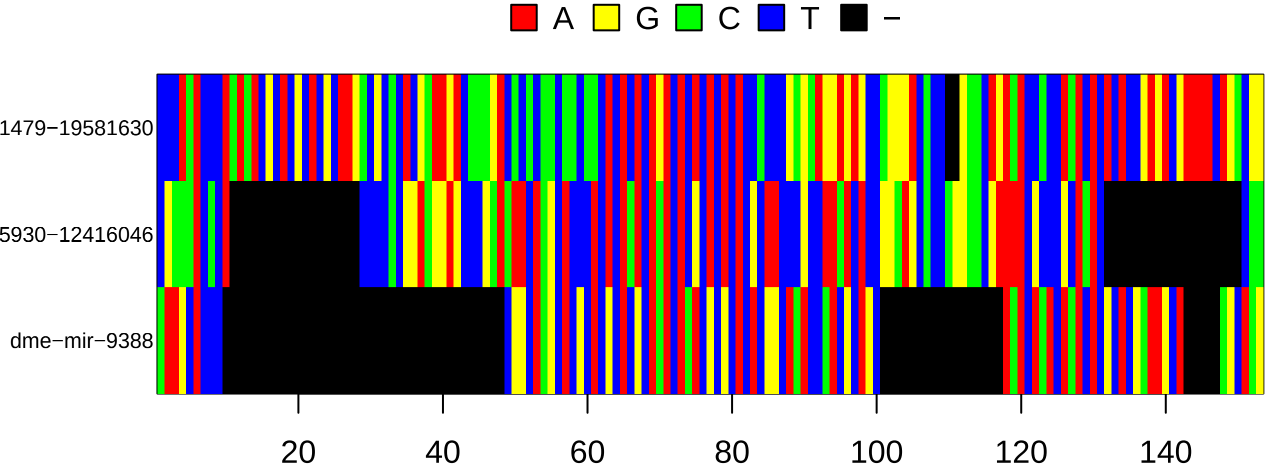
A
G
C
T
−
1479−19581630
5930−12416046
dme−mir−9388
20
40
60
80
100
120
140

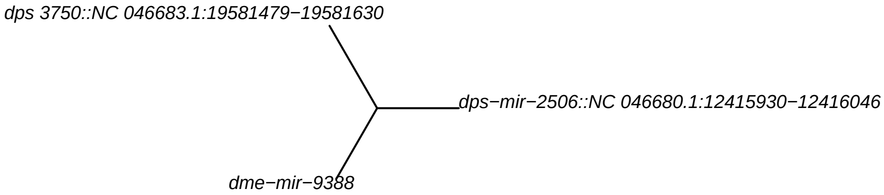
dps 3750::NC 046683.1:19581479−19581630
dps−mir−2506::NC 046680.1:12415930−12416046
dme−mir−9388

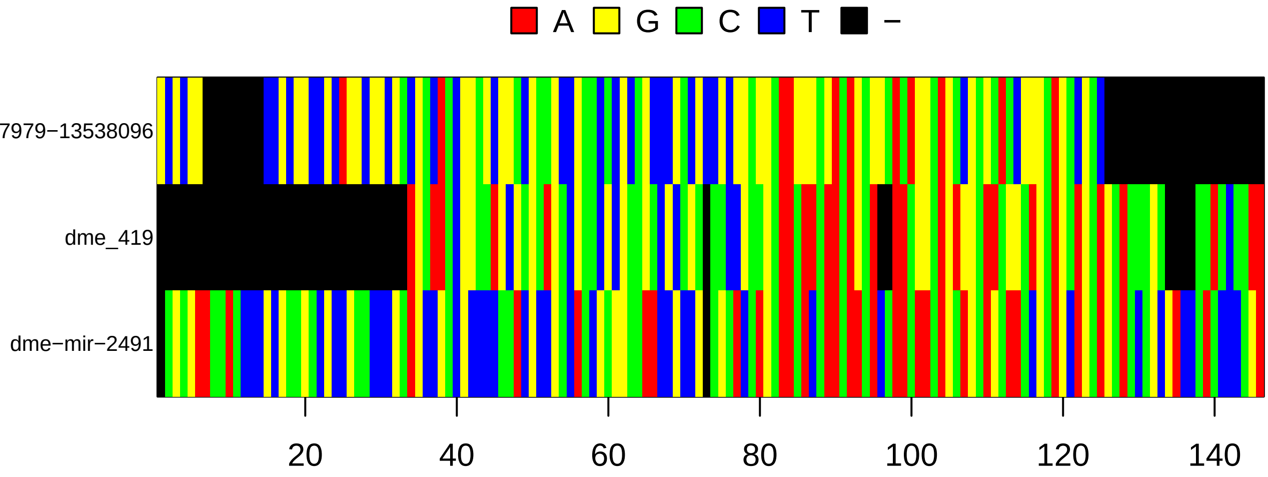
A
G
C
T
−
7979−13538096
dme_419
dme−mir−2491
20
40
60
80
100
120
140

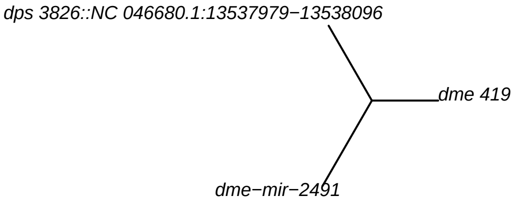
dps 3826::NC 046680.1:13537979−13538096
dme 419
dme−mir−2491

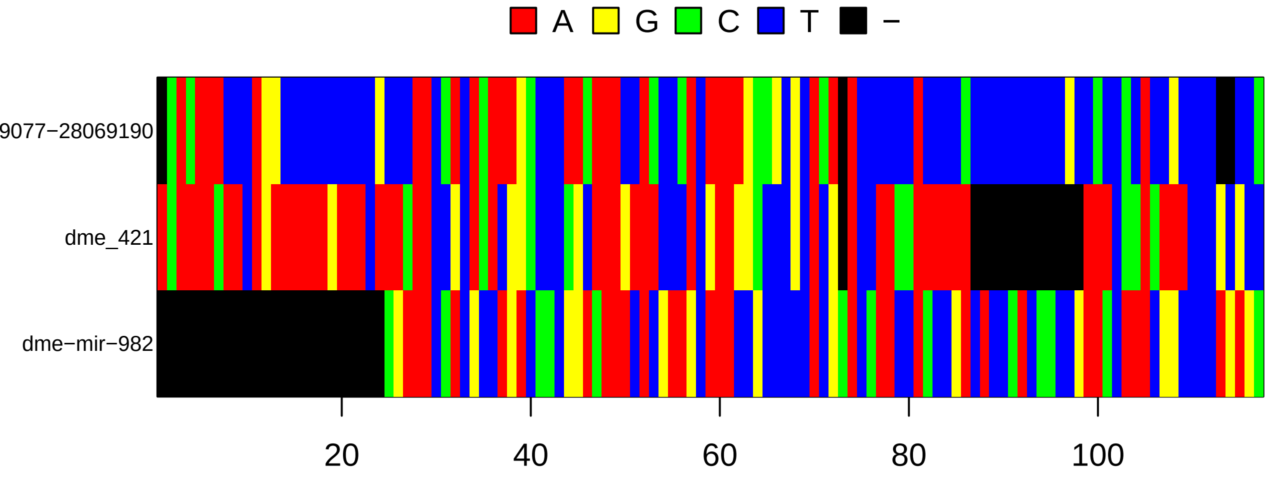

A
G
C
T
−
9077−28069190
dme_421
dme−mir−982
20
40
60
80
100


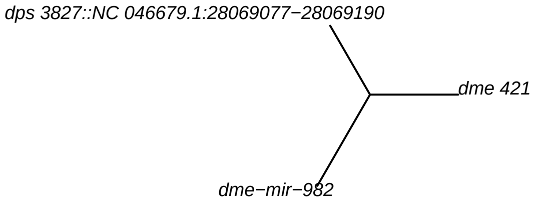

dps 3827::NC 046679.1:28069077−28069190
dme 421
dme−mir−982

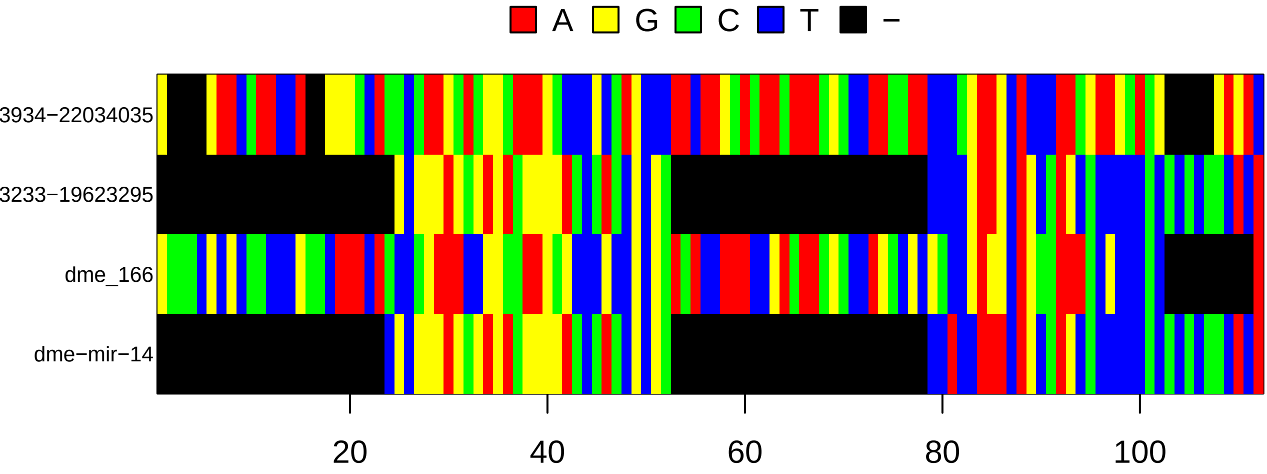
A
G
C
T
−
3934−22034035
3233−19623295
dme_166
dme−mir−14
20
40
60
80
100

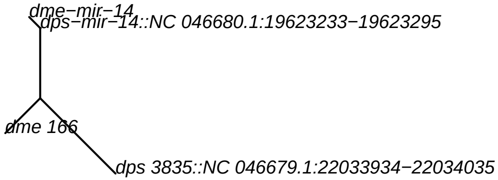
dme−mir−14
dps−mir−14::NC 046680.1:19623233−19623295
dme 166
dps 3835::NC 046679.1:22033934−22034035
