## Supplementary Figure 1 for "The chromosomal distribution of sex-biased microRNAs in *Drosophila* is non-adaptive"

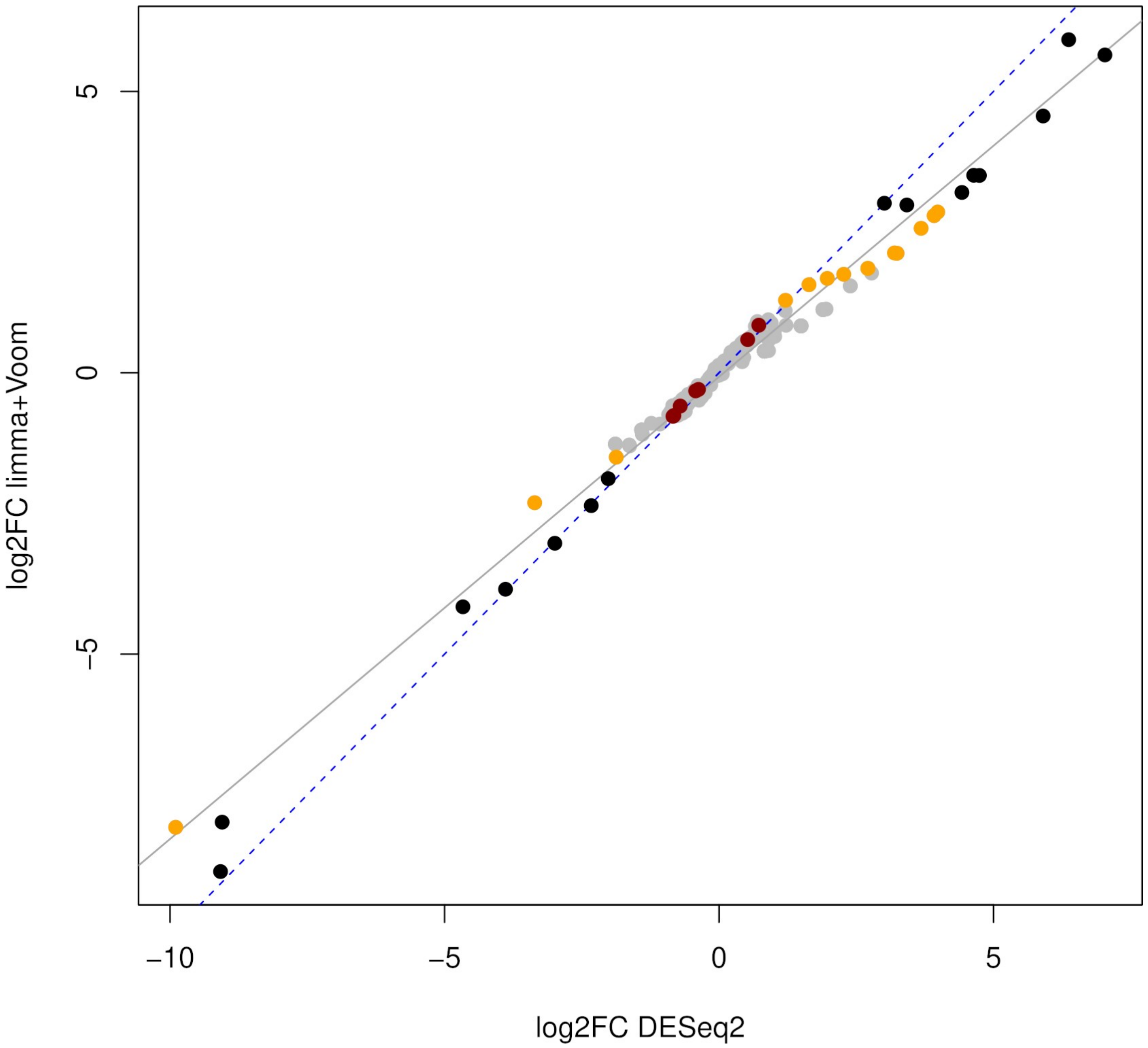


**Supplementary Figure 1.** Comparison between the log2 fold-changes from the differential microRNA expression analysis of the paired samples with DESeq2 and Limma with Voom transformation. Dot color code: black, differentially expressed (DE) according to both methods; orange, DE according to Voom-Limma only; red, DE according to DESeq2 only; grey, not DE with either method.
